# Selective coupling and decoupling prepare distributed brain networks for skilled action

**DOI:** 10.1101/2025.10.19.683309

**Authors:** Stefan M. Lemke, Simone Appaqaq, Jian-Zhong Guo, Adam W. Hantman

## Abstract

The precise and rapid nature of skilled actions has motivated a long-standing theory that preparatory neural states must emerge to enable upcoming actions. This has been best characterized in motor cortex, where neural activity evolves toward an initial state that governs subsequent cortical dynamics. Yet complex actions require coordination across multiple areas beyond motor cortex, and how this distributed network prepares remains unknown. We recorded over 40,000 neurons across the brain as mice produced skilled reach-to-grasp actions, revealing selective coupling of action-informative neurons and decoupling of non-informative neurons over a longer timescale and across a broader network than typically associated with motor preparation. These dynamics predicted upcoming action quality and neural activity on individual trials, and performance was impaired when trials were initiated before coupling and decoupling emerged. Two local field potential rhythms organized these dynamics, providing a target for optogenetic intervention that bidirectionally altered action quality. Our work reveals a distributed preparatory process in which selective coupling and decoupling establish a network state that enables skilled action.

## Introduction

Skilled actions are produced too rapidly to be assembled entirely during execution. This constraint has motivated motor control theories – from motor programs^1,2^ to action representations^3^ – that propose the brain establishes a preparatory neural state before movement that enables skilled action. Yet what defines this state, and how it is established across the brain, remains unclear.

The search for this preparatory state has primarily focused on motor cortex. The first descriptions came in the 1960s, when readiness potentials were identified as negative electroencephalogram shifts over motor cortical areas that develop seconds before movement and covary with upcoming movement features^4,5^. Single-neuron studies starting in the 1970s showed motor cortex firing rates similarly predict movement features during the delay between an instructional cue and movement^6–9^. In the 2000s, large-scale neural recordings formalized the link between neural activity before and during movement, demonstrating that population activity in motor cortex during an instructed-delay period sets the initial state of a dynamical system that governs how neural activity evolves during movement^10,11^.

Yet most skilled actions require coordinated activity across multiple brain areas beyond motor cortex, and how this distributed network prepares for upcoming action is unclear. Recent multi-area studies have shown that additional areas – particularly the thalamus and cerebellum – provide critical inputs that shape the cortical preparatory state^12,13^. However, whether these areas primarily support the cortical state or participate in establishing a more distributed pre-movement state remains an open question.

Skilled reaching and grasping is a complex, ethologically-relevant behavior that exemplifies distributed control, as disruptions to cerebral cortex^14–18^, thalamus^19^, basal ganglia^14,16^, or cerebellum^20–22^ all impair performance. To investigate how these areas prepare to jointly control skilled action, we recorded over 40,000 neurons, simultaneously monitoring frontal and motor cortical areas, striatum, thalamus, and cerebellum as mice produced skilled reach-to-grasp actions. We first identified neurons that encoded information about movement kinematics, revealing a distributed network of action-informative neurons across the brain. We then examined over 6,000,000 cross-area neuron pair interactions and found that, in the ∼10 seconds preceding action production, action-informative neurons across areas coupled while non-informative neurons decoupled. The magnitude of coupling and decoupling predicted the consistency of upcoming neural activity and action quality on a trial-by-trial basis, and disrupting these dynamics by initiating trials early impaired performance. At the population level, two coordinated local field potential rhythms reflected these coupling and decoupling dynamics across the brain.

Optogenetic enhancement or disruption of these rhythms bidirectionally altered action quality, providing causal evidence for their role in skilled action production and highlighting a potential interventional target to enhance skilled action. Our work reveals that the selective coupling of action-informative neurons and decoupling of non-informative neurons establishes a distributed neural state that prepares the brain for upcoming skilled action.

## Results

### Action information is widely distributed across the brain

To study how distributed brain networks prepare for skilled action, we first identified the multi-area network that jointly controls reach-to-grasp actions (**Figure 1a**). Mice achieved skilled reach-to-grasp performance in ten days of training, reflected in expert success rates (∼75-90%), stable trial-to-trial kinematics, and consistent reaction times (**Supp. Figure 1**). We simultaneously recorded from five interconnected brain areas during skilled performance (**Supp. Figure 2**), isolating 43,059 neurons across: (1) frontal cortex (FC) targeting the rostral forelimb area^23^, (2) primary motor cortex (M1) targeting the caudal forelimb area^23^, (3) dorsolateral striatum (DLS), the main basal ganglia target of motor cortex^24,25^, (4) ventroanterior/ventrolateral motor thalamus (mTH), a nucleus receiving basal ganglia and cerebellar input^26^, and (5) the interposed nucleus of the deep cerebellar nuclei (DCN), a cerebellar output critical for skilled reaching^20–22^.

**Figure 1:**
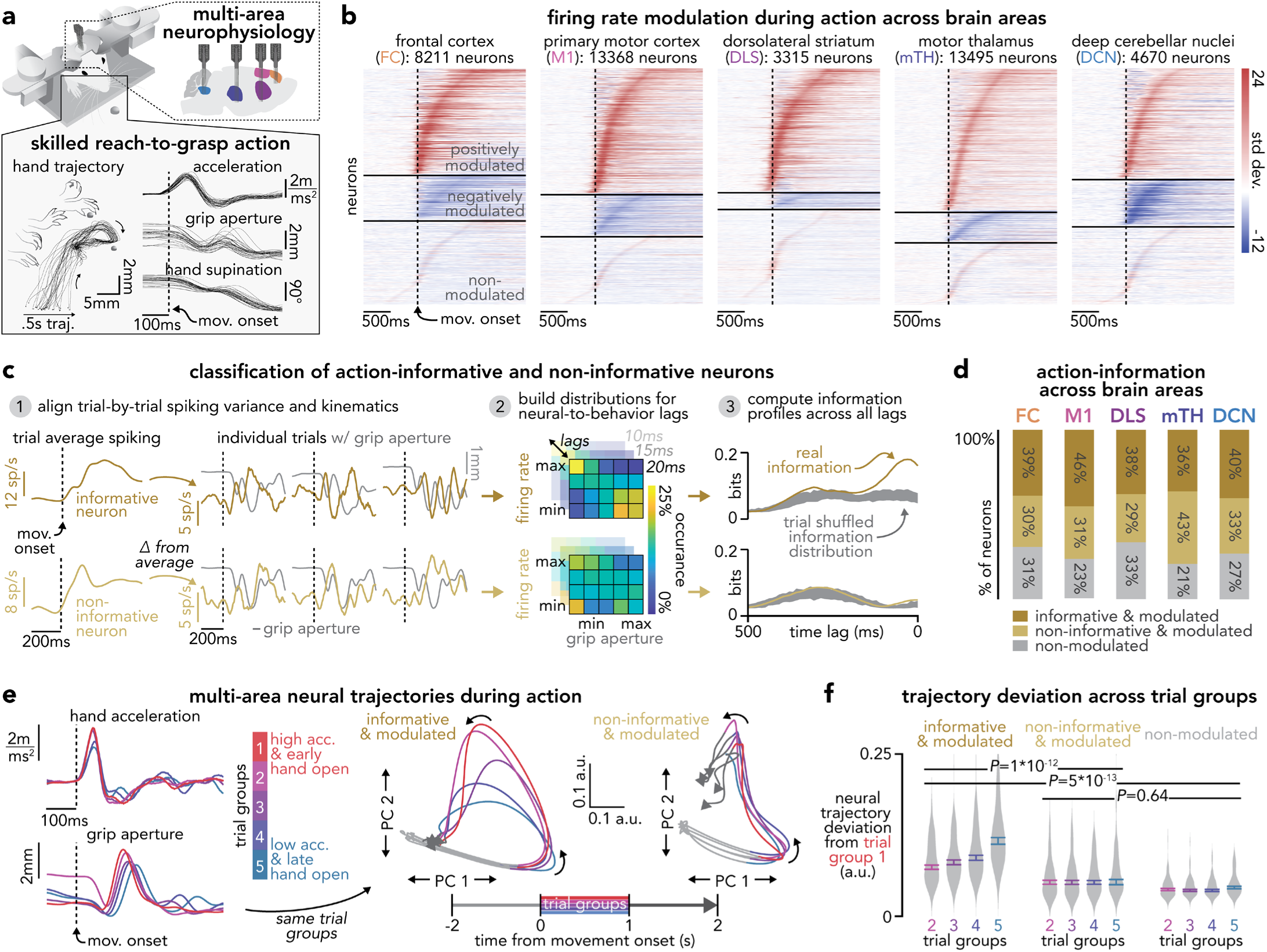
Action information is widely distributed across the brain. **(a)** Schematic of the reach-to-grasp action capturing high-dimensional movement kinematics and simultaneous multi-area neurophysiology. **(b)** Cascade plots of normalized trial-averaged single-neuron firing rates during action production aligned to movement onset for each area, grouped into positively modulated, negatively modulated, and non-modulated neurons (FC: *n* = 8211 neurons from 54 recording sessions from 6 animals, M1: *n* = 13368 neurons, 138 sessions, 17 animals, DLS: *n* = 3315 neurons, 129 sessions, 17 animals, mTH: *n* = 13495 neurons, 95 sessions, 12 animals, DCN: *n* = 4670 neurons, 114 sessions, 16 animals). **(c)** Schematic of action-informative and non-informative neuron classification: (1) trial-by-trial spiking variability was isolated by subtracting median spiking across trials from each trial, (2) Shannon information was computed between this variability and movement kinematics across a range of neural-to-behavior lags, (3) observed information was compared to a trial-shuffled null distribution to identify neurons with significant information. **(d)** Proportion of action-informative modulated, non-informative modulated, and non-modulated neurons in each area (number of neurons matches panel b). **(e)** Left: mean hand acceleration and grip aperture during reaching for five groups of trials from example session, separated based on acceleration magnitude and timing of hand open. Right: mean neural trajectories (top two PCs) from action-informative modulated neurons and non-informative modulated neurons, separated into the same five trial groups based on movement kinematics, from example session. **(f)** Neural trajectory deviation from the top kinematic quintile across trial groups for each neuron population (violin plots in gray show session-level distribution, mean ± s.e.m. across sessions overlaid) The change in deviation between the highest and lowest quintiles was compared across neuron populations using two-sided Wilcoxon signed-rank tests (action-informative/non-informative: *P*<0.001; action-informative/non-modulated: *P*<0.001; non-informative/non-modulated: *P*=0.64, *n* = 90 sessions across 13 animals meeting > 50 neuron criteria).

Neural activity across all areas was robustly modulated during action (**Figure 1b, Supp. Figure 3**). Simultaneous recordings revealed subtle differences in modulation timing: DLS and FC showed the first increases in firing rate relative to movement onset, and DCN the first decrease (**Supp. Figure 3**). However, modulation does not necessarily reflect control of ongoing movement kinematics and could instead reflect general task engagement, cue processing, reward anticipation, or trial-invariant movement signals. To specifically identify neurons whose activity encodes detailed reach-to-grasp kinematics on each trial, rather than simply responding to the task in a stereotyped manner, we computed Shannon information^27^ between each neuron’s trial-to-trial spiking variability and upcoming kinematics (acceleration and grip aperture) at short latency (<500 ms) (**Figure 1c**). A subset of modulated neurons in each area encoded significant kinematic information, which we classified as *action-informative*. The remaining neurons, classified as *non-informative*, lacked this information but may encode other task-related features (**Figure 1d**). Information timing in action-informative neurons mirrored modulation timing, with FC, DLS, and DCN encoding information at the longest neural-to-behavior lags (**Supp. Figure 3**). This revealed a distributed action-informative network that spanned cortex, striatum, thalamus, and cerebellum.

This distributed action-informative network was also evident in population dynamics, as principal component analysis (PCA) trajectories from action-informative neurons tracked trial-by-trial kinematic variation (**Figure 1e,f**). In contrast, trajectories from non-informative or non-modulated neurons did not reflect kinematic differences across trials (**Figure 1e,f**). Neurons contributing to the top principal components were distributed across brain areas, indicating that what neurons encode, rather than their anatomical location, determines whether they share similar dynamics (**Supp. Figure 4**). Given the distributed nature of action-informative neurons across the brain, we next investigated how this multi-area network prepares for skilled action.

### Selective network coupling and decoupling precede skilled action

We examined pre-movement network interactions by measuring spike-timing relationships between 6,658,933 cross-area neuron pairs. To isolate interactions relevant to the upcoming action, rather than stereotyped dynamics occurring on every trial, we computed cross-correlations on median-subtracted spiking activity (to remove trial-invariant spiking activity) as cosine similarity (to normalize across firing rates). This captures whether one neuron’s activity predicts another’s on a trial-by-trial basis. Cross-correlations were computed separately within action-informative and non-informative neuron pairs using a sliding 500 ms window, providing a dynamic readout of interaction (**Figure 2a**). Each cross-area neuron pair was assigned a putative directionality – e.g., DCN to mTH or mTH to DCN – based on the position of the average cross-correlation peak, indicating which area’s spikes led or lagged the other.

**Figure 2:**
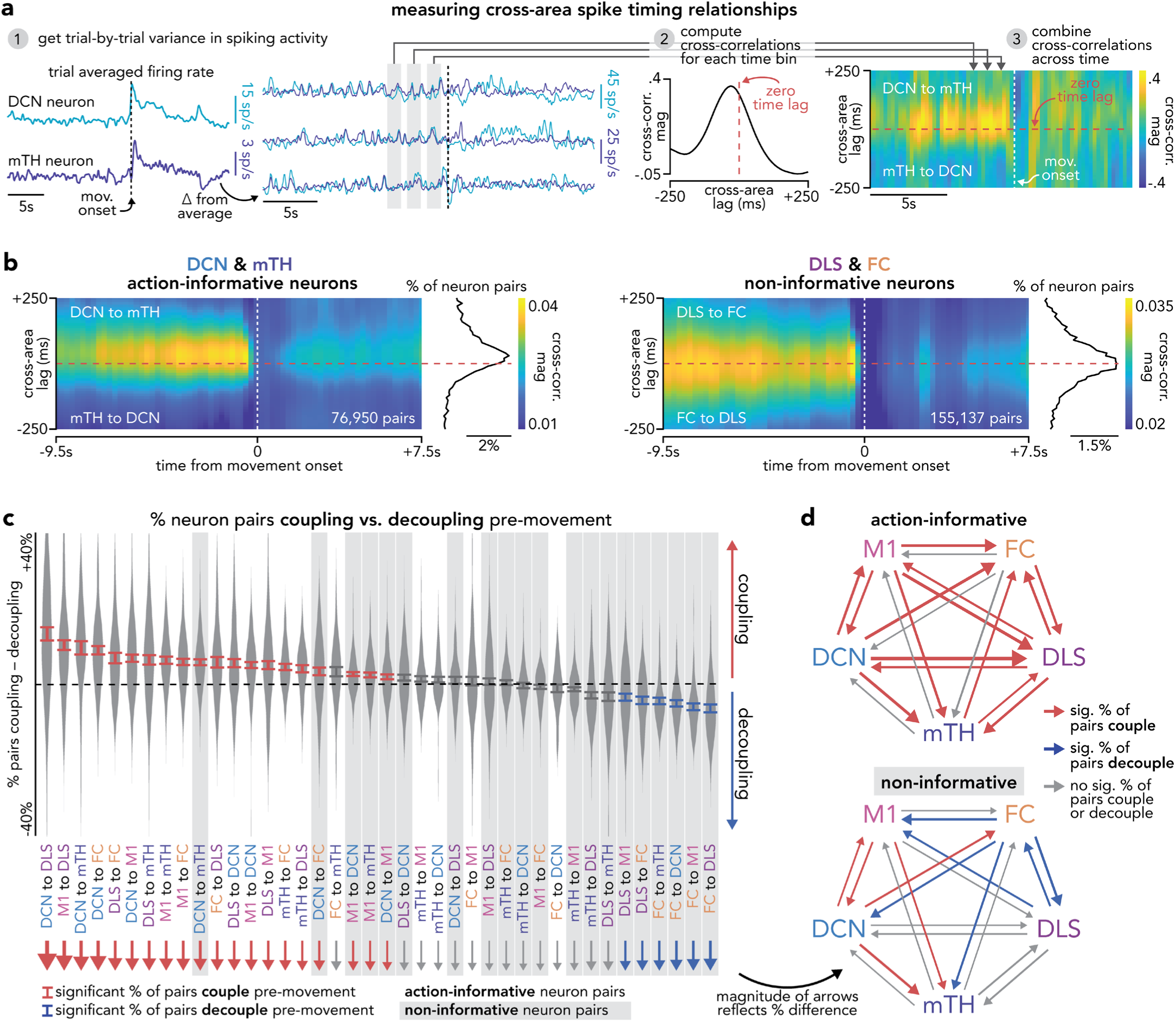
Selective network coupling and decoupling precede skilled action. **(a)** Schematic of cross-correlation computation: (1) trial-by-trial spiking variability was isolated by subtracting median spiking across trials from each trial, (2) cross-correlations between cross-area neuron pairs were computed as cosine similarity within sliding 500 ms bins, (3) cross-correlations were combined to get a dynamic readout of coupling and decoupling. **(b)** Cross-correlation magnitude over time relative to movement onset and across cross-area lags for action-informative DCN-mTH neuron pairs (left, *n* = 76950 neuron pairs from 72 recording sessions from 11 animals) and non-informative DLS-FC neuron pairs (right, *n* = 155137 pairs, 51 recording sessions, 6 animals). Heatmaps show correlation magnitude across time and cross-area time lag, and associated histograms show the distribution of peak cross-correlation lags. **(c)** Difference in percentage of cross-area neuron pairs (in each direction) that increased (coupled) versus decreased (decoupled) cross-correlation magnitude between early (−10 to −6.5 s) and late (−3.75 to −0.5 s) pre-movement periods. Direction of each pair was assigned based on cross-area time lag of cross-correlation peak. Statistical significance was assessed separately within each directional population. Red and blue bars indicate a percentage difference that is significantly above or below zero. Gray shaded background bars indicate distributions from non-informative neuron pairs (*P*<0.05 with Bonferroni correction, one-sample *t*-test). Detailed statistics are in Supp. Table 1. **(d)** Network diagrams summarizing significant changes in percentage of cross-area neuron pairs (for each direction) that couple and decouple during the pre-movement period. Arrows indicate interaction direction and line width reflects magnitude of change. Red = significant increase; blue = significant decrease. Separate diagrams shown for action-informative and non-informative neuron pairs.

We found two dominant changes in cross-area spike-timing consistency across the brain during the 10 s before movement: an increase (coupling) or decrease (decoupling) in spike-timing consistency (**Figure 2b**). We quantified coupling and decoupling between each pair of areas (in each direction) as the proportion of cross-area neuron pairs that increased or decreased in cross-correlation peak magnitude during the 10 s before movement. Across the brain, action-informative neurons primarily coupled, while non-informative neurons decoupled (**Figure 2c, Supp. Table 1**). This pattern reflected action encoding rather than modulation sign, as both positively and negatively modulated neurons coupled and decoupled in similar proportions (**Supp. Figure 5**). Network maps of significant coupling and decoupling revealed a posterior bias for coupling and an anterior bias for decoupling (**Figure 2d**). The most prominent coupling originated from DCN to cerebral regions, accounting for 4 of the 6 most coupled directions, while decoupling was dominated by FC-leading interactions, which accounted for all 4 of the most decoupled directions. In comparison to coupling and decoupling, firing rate changes were localized to specific brain areas and emerged later, closer to movement, consistent with previous work^10,28^ (**Supp. Figure 6**). Together, the selective coupling of action-informative neurons and decoupling of non-informative neurons established a distributed preparatory network state before skilled action.

### Pre-movement coupling and decoupling predict action quality and neural dynamics

To examine how pre-movement coupling and decoupling impact skilled actions, we examined whether these dynamics predicted upcoming neural activity and action quality on a trial-by-trial basis. For each cross-area pair, we measured coupling and decoupling on each trial by averaging the change in cross-correlation peak magnitude during the 10 s pre-movement, across all action-informative neuron pairs that coupled or all non-informative pairs that decoupled. Action quality was quantified as the similarity of each trial’s hand trajectory to the average trajectory of successful trials in that session. Stronger coupling among action-informative pairs predicted higher action quality across the brain (**Figure 3a&c, Supp. Table 2**). In parallel, stronger decoupling among non-informative pairs predicted higher action quality, primarily between anterior brain areas such as FC, M1, and DLS, where we observed the strongest decoupling (**Figure 3b&c, Supp. Table 2**). In contrast, pre-movement firing rate changes were weaker predictors of action quality (**Supp. Figure 7**). Coupling and decoupling also predicted the consistency of neural dynamics during action, with stronger coupling and decoupling associated with greater similarity between each trial’s neural trajectory and the average trajectory on successful trials (**Figure 3c, Supp. Table 2**). This suggests that pre-movement coupling of action-informative neurons and decoupling of non-informative neurons enables the subsequent production of consistent neural dynamics and skilled actions.

**Figure 3:**
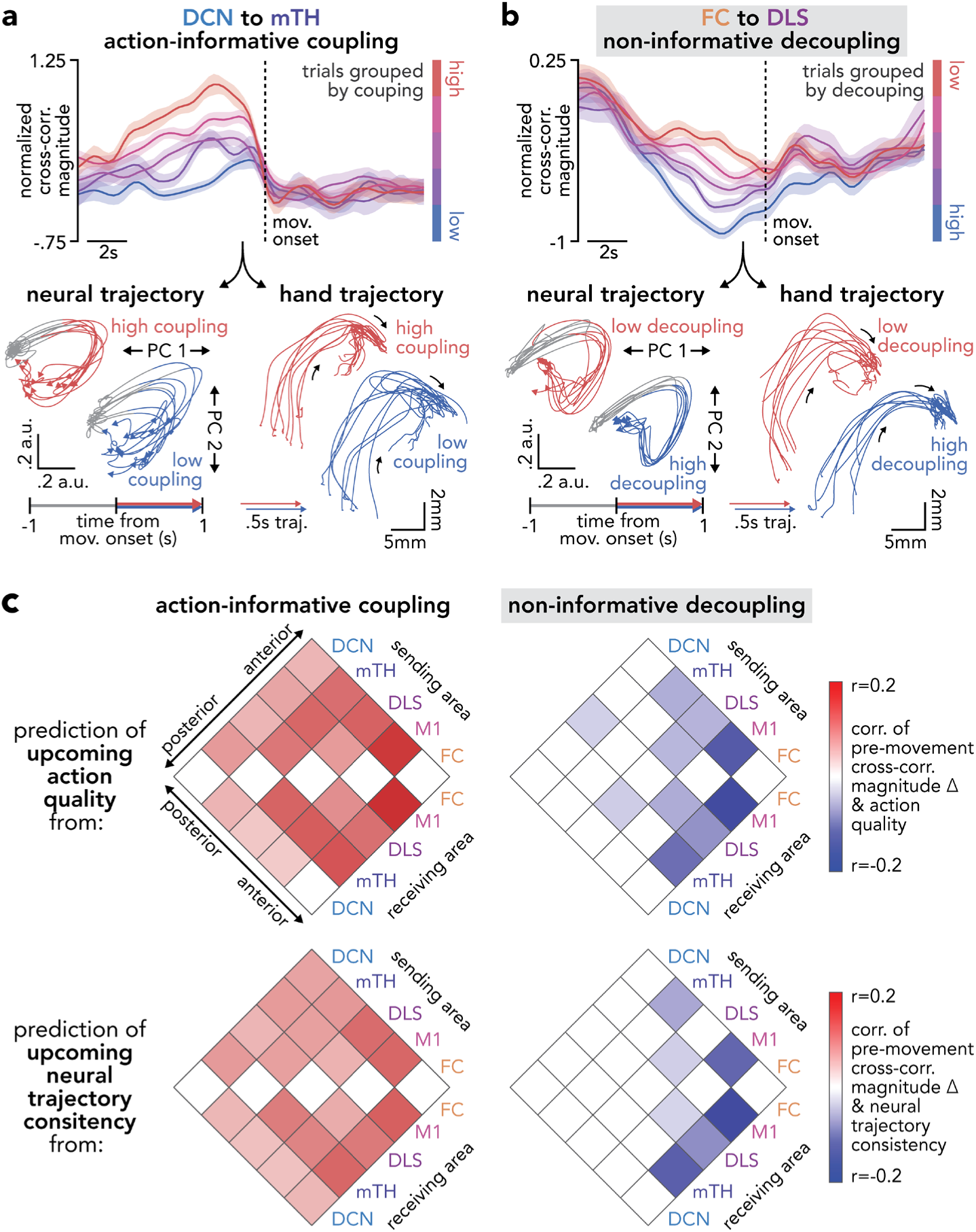
Pre-movement coupling and decoupling predict action quality and neural dynamics. **(a)** Cross-correlation magnitude over time for DCN-to-mTH action-informative neuron pairs that coupled, separated into five trial groups based on pre-movement coupling magnitude (top, line width denotes mean ± s.e.m. across trials in each group, *n* = 273 trials in 5 subgroups from five sessions in one example animal). Corresponding neural trajectories and hand trajectory kinematics are shown for the lowest and highest coupling groups (bottom). **(b)** Same as **(a)** for non-informative FC-to-DLS neuron pairs grouped by pre-movement decoupling magnitude (*n* = 390 trials in 5 subgroups from five sessions in one example animal). **(c)** Matrix of correlations relating trial-by-trial changes in pre-movement cross-correlation magnitude within action-informative neuron pairs that coupled (left) and non-informative neuron pairs that decoupled (right) to upcoming action quality (top) and neural trajectory consistency (bottom). Color indicates significant correlation at *P*<0.05, Bonferroni-corrected (Spearman correlation). Detailed statistics are in Supp. Table 2.

### Disrupting coupling and decoupling impairs action quality and neural dynamics

To test the necessity of these pre-movement dynamics for skilled action, we manipulated the inter-trial interval to determine if starting trials before typical coupling and decoupling emerged would impair performance. After training mice with a consistent 30 s inter-trial interval, we introduced trials at 20 s (early), 30 s (regular), or 40 s (late) intervals. Early trial initiation significantly reduced both action-informative coupling and non-informative decoupling compared to regular and late trials (**Figure 4a&b**), although reaction times were similar (early: 149.2 ± 3.8 ms; regular: 144.2 ± 4.2 ms; late: 141.6 ± 5.4 ms; early/regular: *P*=0.29; regular/late: *P*=0.59, mean ± s.e.m.; two-sided Wilcoxon signed-rank tests, *n* = 17 sessions). Consistent with pre-movement coupling and decoupling enabling skilled action, early trials had significantly reduced action quality and neural trajectory consistency compared to regular and late trials (**Figure 4a&c**), highlighting the importance of pre-movement coupling and decoupling dynamics for skilled action.

**Figure 4:**
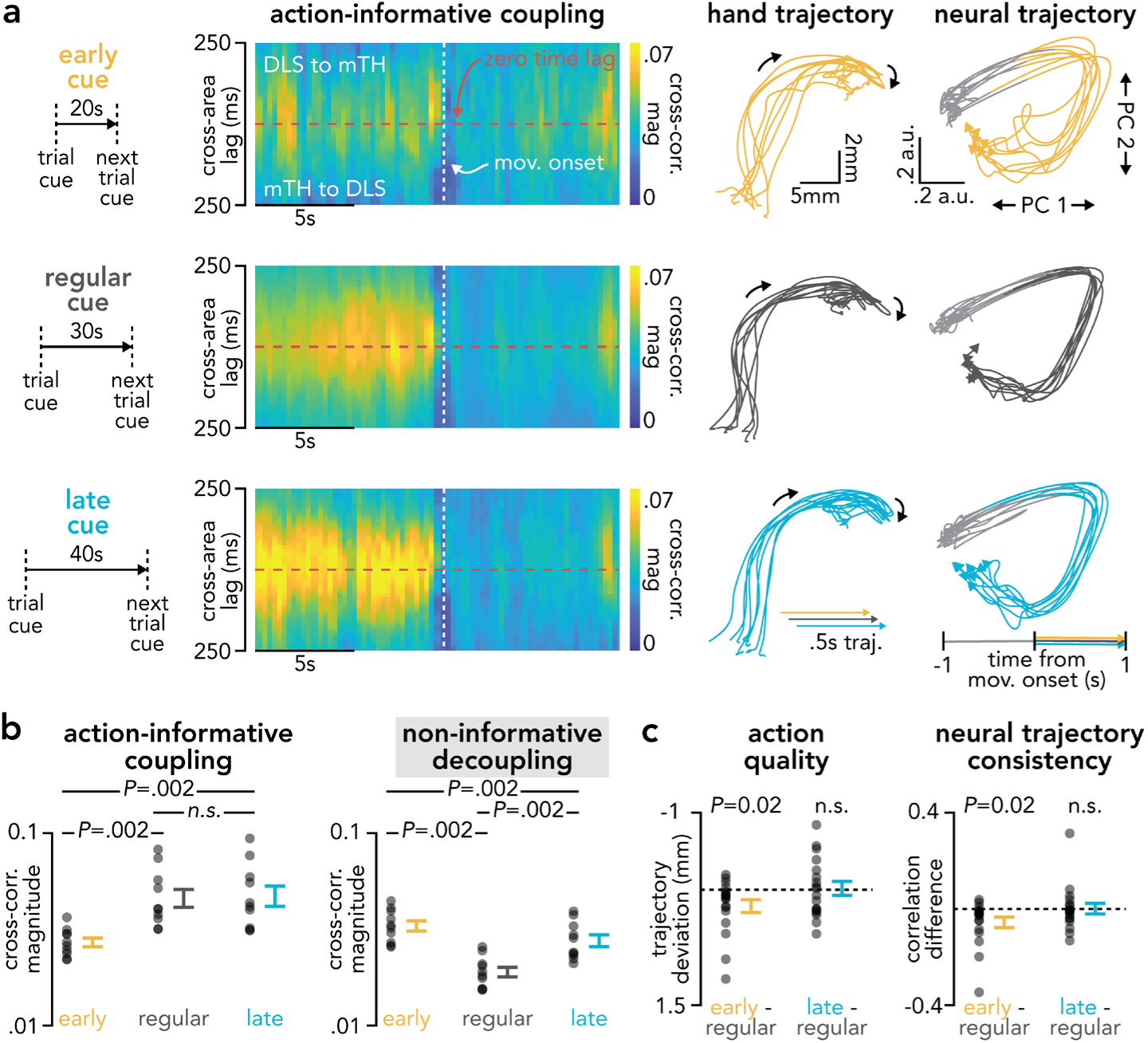
Disrupting coupling and decoupling impairs action quality and neural dynamics. **(a)** Schematic of inter-trial interval manipulation and cross-correlation magnitude over time relative to movement onset and across cross-area time lags for action-informative DCN-mTH neuron pairs during early, regular, and late cue trials (*n* = 17 recording sessions from 4 animals). Example hand trajectory kinematics and neural trajectories are shown for each condition. **(b)** Comparison of average pre-movement cross-correlation magnitude within each cross-area pair during early, regular, and late cue trials for action-informative neuron pairs that couple (left) and non-informative pairs that decouple (right). Two-sided Wilcoxon signed-rank test (*n* = 10 cross-area pairs). Action-informative: early/regular: *P*=0.002; early/late: *P*=0.002; regular/late: *P*=0.19. Non-informative: early/regular: *P*=0.002; early/late: *P*=0.002; regular/late: *P*=0.002. **(c)** Comparison of action quality (left) and neural trajectory consistency (right) for early and late cue trials relative to regular trials (two-sided Wilcoxon signed-rank test, *n* = 17 sessions from 4 animals). Action quality: early: *P*=0.02; late: *P*=0.95. Neural trajectory consistency: early: *P*=0.007; late: *P*=0.45.

### Coordinated brain rhythms organize pre-movement coupling and decoupling

The distributed nature of pre-movement coupling and decoupling across the brain suggested a coordinated organizing process. Local field potential (LFP) rhythms were a natural candidate: cross-area LFP coherence can dynamically gate inter-area communication^29^ and pre-movement changes in LFP rhythms, such as beta oscillations^30^, are well-established. Yet, whether these population-level rhythms organize specific distributed networks of neurons is unclear. We computed changes in LFP coherence across all areas during the 10 s preceding movement onset and found two LFP rhythms with opposing pre-movement dynamics: an emerging delta-band (δ, 4 Hz peak) coherence, most prominent between posterior areas such as DCN and mTH, and a decreasing beta-band (β, 12 Hz peak) coherence, most prominent between anterior areas such as FC and M1 (**Figure 5a&b**). The posterior/anterior bias of δ increases and β decreases mirrored neuron-level coupling and decoupling patterns (**Figure 5b**, **Figure 2d**). Pre-movement changes in LFP power showed a similar pattern, with increased δ power localized to DCN and decreased β power observed across cerebral regions (**Supp. Figure 8a**). The spatial correspondence between LFP dynamics and neuron-level coupling and decoupling suggests oscillatory coherence as a potential organizing mechanism for neuron-level dynamics.

**Figure 5:**
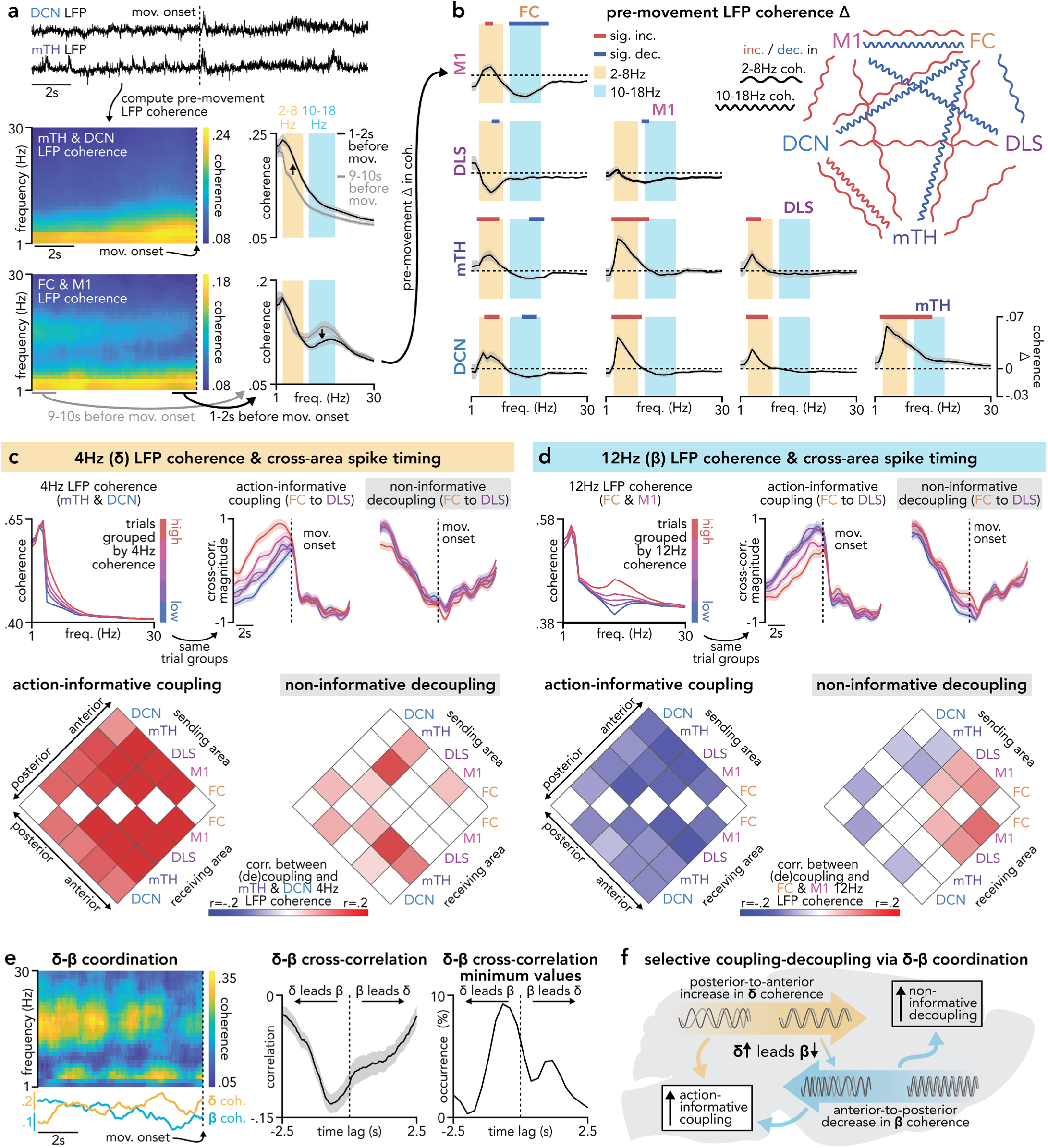
Coordinated brain rhythms organize pre-movement coupling and decoupling. **(a)** Example computation of pre-movement LFP coherence between DCN-mTH (top) and FC-M1 (bottom). Raw LFP traces aligned to movement onset and corresponding coherence spectrogram with frequency-specific coherence spectra from early (gray) and late (black) pre-movement periods, highlighting 2-8 Hz (yellow background) and 10-18 Hz (blue background) frequency bands. Line width denotes mean ± s.e.m. across sessions (mTH-DCN: *n* = 100 sessions across 12 animals; FC-M1: *n* = 55 sessions across 6 animals). **(b)** Changes in pre-movement LFP coherence between early (9-10 s before movement) and late (1-2 s before movement) pre-movement periods across all recorded area pairs. Coherence spectra show significant increases (red bars) or decreases (blue bars), highlighting 2-8 Hz (yellow background) and 10-18 Hz (blue background) frequency bands (Wilcoxon signed-rank test, Bonferroni-corrected, *P*<0.05). Network schematic summarizes significant increases (red) or decreases (blue) in cross-area coherence within 2-8 Hz (low-frequency squiggly lines) or 10-18 Hz (high-frequency squiggly lines). FC-M1: *n* = 55 sessions across 6 animals; FC-DLS: *n* = 55 sessions across 6 animals; FC-mTH: *n* = 52 sessions across 6 animals; FC-DCN: *n* = 54 sessions across 6 animals; M1-DLS: *n* = 155 sessions across 17 animals; M1-mTH: *n* = 100 sessions across 12 animals; M1-DCN: *n* = 145 sessions across 16 animals; DLS-mTH: *n* = 100 sessions across 12 animals; DLS-DCN: *n* = 145 sessions across 16 animals; mTH-DCN: *n* = 100 sessions across 12 animals. **(c)** Relationship between pre-movement 4 Hz (delta, δ) LFP coherence and cross-area spike coupling and decoupling. Top: mTH-DCN coherence spectra for five trial groups based on δ coherence magnitude, with corresponding average FC to DLS action-informative coupling and non-informative decoupling over time shown for the same trial groups (*n =* 3601 trials from 51 sessions across 6 animals). Bottom, matrices showing correlations between pre-movement mTH-DCN δ coherence and action-informative coupling (left) or non-informative decoupling (right) across all sending/receiving area pairs. Color indicates significant correlation at *P*<0.05, Bonferroni-corrected (Spearman correlation). Detailed statistics are in Supp. Table 3. **(d)** Same as **(c)** for pre-movement 12 Hz (beta, β) LFP coherence between FC and M1 (*n =* 2976 trials from 43 sessions across 6 animals). Detailed statistics are in Supp. Table 3. **(e)** δ-β coordination during the pre-movement period. Left: example session LFP coherence spectrogram and corresponding δ and β coherence time courses. Middle: average cross-correlation between δ and β coherence across all cross-area pairs, for sessions with pre-movement δ coherence increases and β coherence decreases, computed separately for each cross-area pair within each animal (line width represents mean ± s.e.m. across cross-area pairs pooled across animals, *n* = 92 cross-area pair observations across 12 animals). Right: distribution of lag values at which the cross-correlation reached its minimum (strongest anti-correlation). **(f)** Schematic summarizing how coordinated δ-β rhythms selectively organize action-informative coupling and non-informative decoupling across distributed brain networks.

LFP coherence and coupling/decoupling were also closely aligned on a trial-by-trial basis. We measured trial-by-trial coherence between the areas with the largest pre-movement changes: mTH-DCN for δ coherence and FC-M1 for β coherence. On each trial, we compared LFP coherence during the 5 s preceding movement onset to coupling among action-informative pairs and decoupling among non-informative pairs. Across the brain, δ coherence was positively correlated with action-informative coupling (**Figure 5c, Supp. Table 3**). Consistent with this relationship, action-informative neurons were preferentially phase-locked to δ-band LFP signals pre-movement (**Supp. Figure 9**), suggesting that the consistency of cross-area spike-timing relationships may be supported by this emerging δ coherence. In contrast, β coherence showed a bidirectional relationship: negatively correlated with action-informative coupling across the brain and positively correlated with non-informative decoupling between anterior areas (**Figure 5d, Supp. Table 3**). This bidirectional relationship between anterior areas mirrored the unique combination of coupling and decoupling observed in these areas (**Figure 2d**) and is consistent with work in primates proposing that β oscillations prevent the initiation of movement-generating neural activity^31^. Neural entrainment to β-band LFP signals was substantially weaker than to δ-band LFP (action-informative: 9.7 ± 2.2% phase locked neurons, non-informative: 4.1 ± 2.2%, mean ± s.e.m. across 5 areas). Since both δ and β coherence were correlated with trial-by-trial coupling and decoupling, we next examined how these rhythms interacted.

As β oscillations are transient bursts that appear sustained only when averaged across trials^32^, we measured δ and β interaction by examining their temporal relationships on individual sessions (**Figure 5e**). This revealed a consistent temporal coordination with increases in δ coherence preceding decreases in β coherence before movement (**Figure 5e**). This same coordination also occurred in pre-movement LFP power (**Supp. Figure 8b**). Together, these results suggest a model bridging oscillatory coherence to spike-timing dynamics of distributed neural populations: temporally coordinated increases in posterior δ coherence and subsequent decreases in anterior β coherence orchestrate action-informative coupling and non-informative decoupling among cross-area neuron pairs to prepare the relevant distributed network for skilled action (**Figure 5f**).

### Pupil dynamics reflect network coupling and decoupling

The multi-second timescale of these network dynamics is consistent with neuromodulatory influence. As pupil size can reflect neuromodulatory tone^33^, we measured pupil dynamics to explore this possibility. Pupil size gradually increased in the 10 s before movement (**Supp. Figure 10a**). On a trial-by-trial basis, pupil size was significantly positively correlated with δ coherence and negatively correlated with β coherence (**Supp. Figure 10b, Supp. Table 4**). Pupil size was also significantly positively correlated with action-informative coupling across the brain and non-informative decoupling in anterior areas (**Supp. Figure 10c, Supp. Table 4**). This identifies pupil dynamics as a readout of network coupling and decoupling and suggests a role for neuromodulators such as acetylcholine and norepinephrine in organizing neural populations before action^33^.

### Manipulating cross-area rhythms bidirectionally impacts action quality

We next asked if manipulating cross-area coupling and decoupling before movement could impact skilled action. While selectively manipulating action-informative neurons across the brain is not currently possible, an alternative is to design interventions that target brain rhythms that preferentially entrain these neurons. The relationship we identified between δ/β coherence and coupling/decoupling offered an opportunity to test this approach. Because δ coherence correlated with coupling, preferentially entrained action-informative neurons, and preceded β coherence decreases – which correlated with both coupling and decoupling – we targeted δ as a central organizing rhythm, focusing on DCN, mTH, and M1, where pre-movement δ increases were largest. δ coherence may promote cross-area coupling by ensuring output from DCN reaches mTH at an optimal phase, consistent with theories of oscillatory gating of inter-area communication^29^. If so, altering the relative phase between DCN and mTH should enhance or disrupt this cross-area coordination and, in turn, action quality.

To test this, we optogenetically modulated cortical and cerebellar neural populations in VGAT-ChR2 expressing mice (**Figure 6a**). Two optical fibers targeted lobule simplex of the cerebellar cortex (which projects to the interposed nucleus of the DCN) and M1 (which projects to mTH). We simultaneously recorded neural activity in the DCN and mTH. On each trial, 4 s of 4 Hz sinusoidal light stimulation was delivered to both areas, ending 500 ms before the trial cue. The phase difference between cerebellar and cortical stimulation was systematically shifted across 16 trial types, sampling a full cycle of relative phase differences (**Figure 6a**). Most mTH and DCN neurons were entrained to cortical and cerebellar stimulation, respectively (**Figure 6b**). mTH neurons showed consistent phase locking at a single preferred phase relative to motor cortex stimulation, while DCN neurons exhibited a broader distribution of preferred phases relative to cerebellar stimulation, potentially reflecting greater diversity in cerebellar inputs to DCN compared to cortical inputs to mTH. Consistent with this entrainment, we drove a range of spike-timing relationships across trial types that systematically varied in similarity to the physiological spike-timing relationship measured between DCN and mTH on successful non-laser trials, where DCN spikes led mTH spikes at short latency (∼10-20 ms) (**Figure 6c&d**). This allowed us to test the impact of driving spike-timing at physiological versus non-physiological cross-area timings on upcoming neural activity and action quality. We found a bidirectional impact on behavior: phase differences that produced physiological spike-timing relationships improved action quality, while non-physiological phases disrupted it (**Figure 6e**). We also found that physiological cross-area timing reduced thalamic β power (**Figure 6f**), consistent with the coordination we observed between δ and β coherence, suggesting that δ dynamics may contribute to β suppression. This demonstrates that pre-movement cross-area spike-timing relationships influence upcoming action quality and identifies pre-movement brain rhythms as tractable targets for intervention.

**Figure 6:**
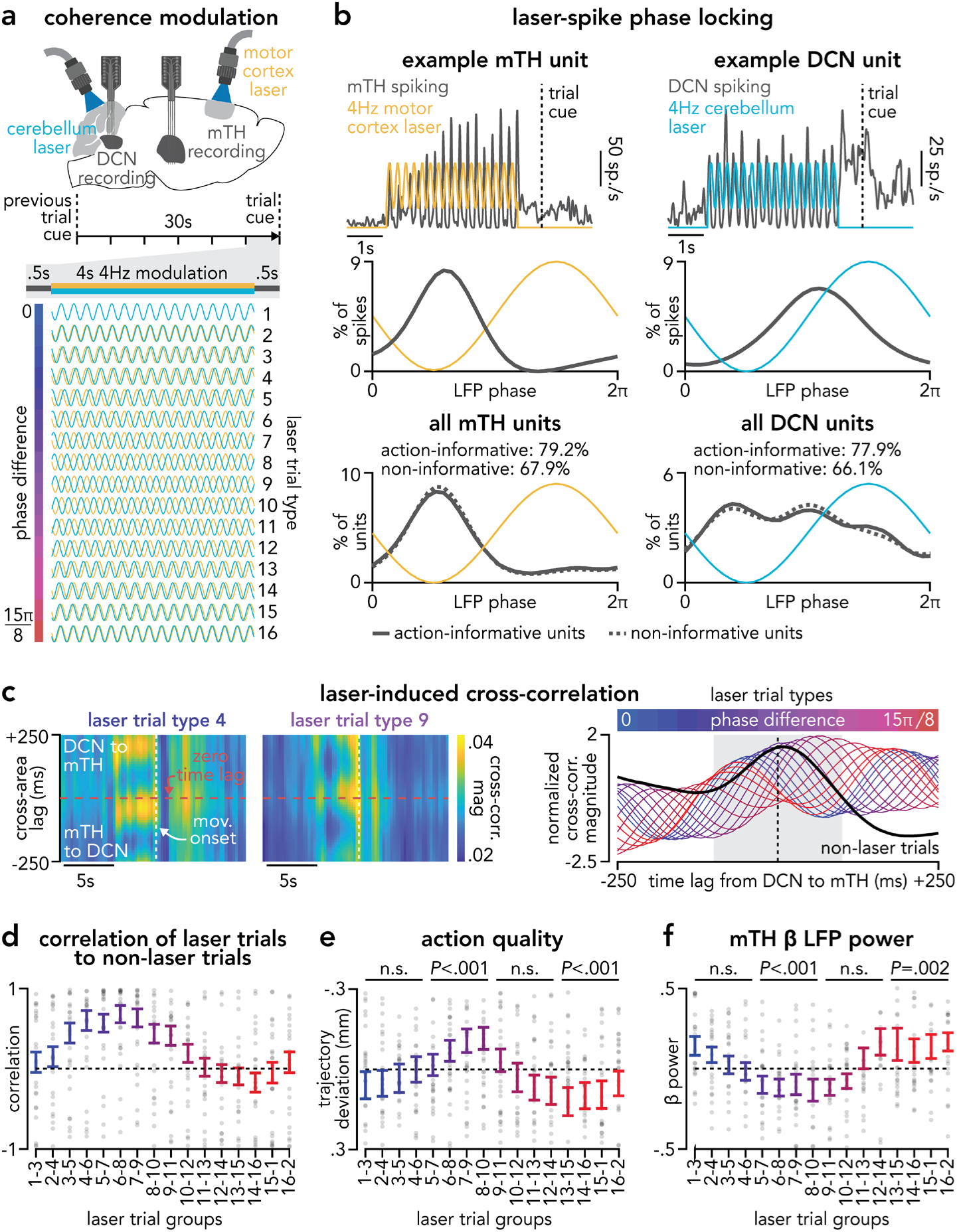
Manipulating cross-area rhythms bidirectionally impacts action quality. **(a)** Schematic of dual-site optogenetic stimulation in VGAT-ChR2 mice. Two optical fibers targeted lobule simplex of the cerebellar cortex and M1, with simultaneous DCN and mTH recordings. On each trial, 4 s of 4 Hz sinusoidal light stimulation was delivered with one of 16 relative phase offsets (0 to 15π/8), ending 500 ms before the trial cue. Each session began and ended with no-laser control trials. **(b)** Phase locking of DCN and mTH neurons to laser stimulation. Top: example mTH (left) and DCN (right) unit spiking during 4 Hz motor cortex and cerebellar stimulation, respectively. Middle: spike phase distribution relative to laser for the example units shown above. Bottom: distribution of preferred phase across all significantly phase-locked mTH and DCN units (DCN neurons: *n* = 512 action-informative (77.9% phase locked), 510 non-informative (66.1% phase locked); mTH neurons: *n* = 3651 action-informative (79.2% phase locked), 3040 non-informative (67.9% phase locked), from 24 sessions across 5 animals). Action-informative (solid) and non-informative (dashed) neurons plotted separately. Percentages indicate the proportion of neurons significantly phase-locked (Rayleigh test, P<0.05, Bonferroni-corrected). **(c)** Cross-area spike-timing relationships during laser stimulation. Left: example cross-correlation magnitude over time relative to movement onset and across cross-area time lags for action-informative DCN-mTH neuron pairs for two example laser trial types (types 4 and 9, *n* = 24 sessions across 5 animals). Right: trial-averaged cross-correlation profiles across time lags for each laser trial type from 24 sessions across 5 animals (types 1-16, blue to red) overlaid on non-laser trials (black). Gray shaded area denotes time lags used to compute correlation in panel **d**. **(d)** Similarity of laser-induced cross-correlation to successful no-laser control trials. Pearson correlation between cross-correlation profiles for sliding groups of 3 consecutive laser trial types (with wrap-around) and non-laser control trials (each gray dot represents one session and error bars represent mean ± s.e.m. across sessions, *n* = 24 sessions across 5 animals). **(e)** Action quality across groups of laser trial types, relative to the median across all 16 types. Lower trajectory deviation indicates better action quality (each gray dot represents one session and error bars represent mean ± s.e.m. across sessions, *n =* 38 sessions across 6 animals, including sessions without neural recording). Statistical significance assessed using two-sided Wilcoxon signed-rank tests vs. zero (groups 1-4: *P*=0.19; groups 5-8: *P*<0.001; groups 9-12: *P*=0.2; groups 13-16: *P*<0.001). **(f)** mTH β LFP power across groups of laser trial types, relative to the median across all 16 types (each gray dot represents one session and error bars represent mean ± s.e.m. across sessions, *n* = 24 sessions across 5 animals). Statistical significance assessed using two-sided Wilcoxon signed-rank tests vs. zero with Bonferroni correction (groups 1-4: *P*=0.02; groups 5-8: *P*<0.001; groups 9-12: *P*=0.8; groups 13-16: *P*=0.003).

## Discussion

### Motor preparation as a distributed network-level process

In the mammalian brain, individual neurons contact hundreds to thousands of local and distant neuron partners, forming highly distributed networks^34–36^. Consistent with this architecture, neural activity across cerebral and cerebellar areas is broadly modulated during purposeful movements^37–40^. To generate a complex action, this distributed activity must be coordinated across the brain to produce appropriate motor output. Here, we reveal a principle by which the brain enables such coordination: before movement begins, neurons carrying information about the upcoming action become increasingly coordinated across the brain, while neurons without such information disengage, establishing a distributed preparatory state that ensures upcoming neural dynamics evolve appropriately to produce skilled action. This state predicts both upcoming action quality and the consistency of neural activity on individual trials, and is necessary for skilled movement: performance is impaired when movements are initiated before this state emerges and optogenetic manipulation of the cross-area rhythms that organize this state bidirectionally alters upcoming action quality. This preparatory state emerges on a longer timescale and across a more distributed network than is typically associated with motor preparation, suggesting it operates alongside a shorter-timescale preparation characterized by firing rate changes in motor cortex.

### Behaviorally relevant neural populations exhibit opposing cross-area dynamics

Understanding how distributed networks jointly control skilled actions requires isolating behaviorally relevant interactions from global cross-area dynamics. We show that distinct neural populations between two areas can simultaneously exhibit opposing forms of interaction: coupling among action-informative neurons and decoupling among non-informative neurons. This is consistent with work demonstrating that task-relevant cross-area dynamics can substantially differ from overall, non-specific measures of neural interaction^41^. These results emphasize the need for high-yield single-neuron-resolution multi-area studies to distinguish diverse forms of interaction that coexist within distributed neural populations. Linking such single-neuron dynamics to emerging analytical tools that identify behaviorally relevant subspaces^42–45^ will help formalize how these distributed networks selectively coordinate to control complex movements. It is possible that selective coupling and decoupling represent a general principle by which the brain organizes distributed networks for goal-directed behavior, integrating the established roles of frontal cortex^46^ and thalamus^47^ in routing and coordinating task-relevant activity during cognitive tasks. The multi-second timescale of this coordination also suggests a role for neuromodulatory systems, consistent with our finding that pupil size correlates with multiple aspects of this preparatory state^33^. How neuromodulatory state, which may reflect pre-movement behavioral state changes such as arousal, and network coupling and decoupling interact remains an important open question.

### Brain rhythms selectively organize cross-area neuron-level dynamics

The widespread coupling and decoupling of neurons across the brain was mirrored at the population level by two coordinated LFP rhythms: an emerging delta (δ, 4 Hz peak) coherence across areas with strongest neuron-level coupling and a decreasing beta (β, 12 Hz peak) coherence across areas with strongest neuron-level decoupling. These opposing dynamics were distributed along a posterior-anterior gradient, with the largest δ coherence increases in posterior areas (e.g., DCN/mTH) and β coherence decreases in anterior areas (e.g., FC/M1). While pre-movement β suppression in cortex is well established^30^, its link to neuron-level dynamics remains unclear. By linking decreases in β coherence to both coupling and decoupling, our results support the idea that β suppression enables the coordination of action-relevant neurons and the concurrent disengagement of neurons that may interfere with the upcoming action^31^. Unexpectedly, this β suppression was preceded by increases in δ coherence, suggesting a hierarchical organization in which posterior δ rhythms contribute to anterior β suppression, in line with established theories of cross-frequency organization^48,49^. The correlation of these rhythms with neuron-level coupling and decoupling on a trial-by-trial basis is consistent with theories of oscillatory gating, in which coherence between areas enables inputs to arrive at a specific phase that either promotes or disrupts consistent spike timing^29^. Together, these results indicate that population-level brain rhythms and single-neuron interactions are expressions of the same emerging preparatory network state.

### Pre-movement brain rhythms as a therapeutic target

While directly targeting action-informative neurons distributed across the brain is not currently feasible, targeting the brain rhythms that organize them is. The pre-movement period offers a natural window for such interventions, when sensory and motor drive are low and network coordination can be monitored and selectively targeted. Our multi-site optogenetic intervention provides proof-of-principle that pre-movement oscillatory dynamics are a tractable target for shaping network interactions, with particular relevance for conditions characterized by atypical cross-area coordination.

## Acknowledgements.

This study was supported by grants from the NICHD/NIH T32HD040127 (S.M.L.), NINDS/NIH F32NS131217 (S.M.L.), NINDS/NIH R01NS118030-01A1 (A.W.H.). The authors thank members of the Hantman laboratory for helpful comments, guidance, and discussion about the manuscript. The authors thank Jeremy D. Cohen for technical assistance and Scott T. Albert and Trisha V. Vaidyanathan for feedback on the manuscript.

## Author contributions

S.M.L., J-Z.G., and A.W.H. conceptualized the project and designed the experiments. S.M.L., S.A., and J-Z.G. performed surgeries and collected behavioral and neural data. S.M.L. analyzed the data. S.M.L. wrote the manuscript with input from A.W.H. A.W.H. and J-Z.G. supervised the project.

## Materials & Correspondence

All inquiries can be sent to A.W.H. and S.M.L. All the source data and code used in this work will be publicly available upon final publication.

---

**Supplemental Figure 1:**
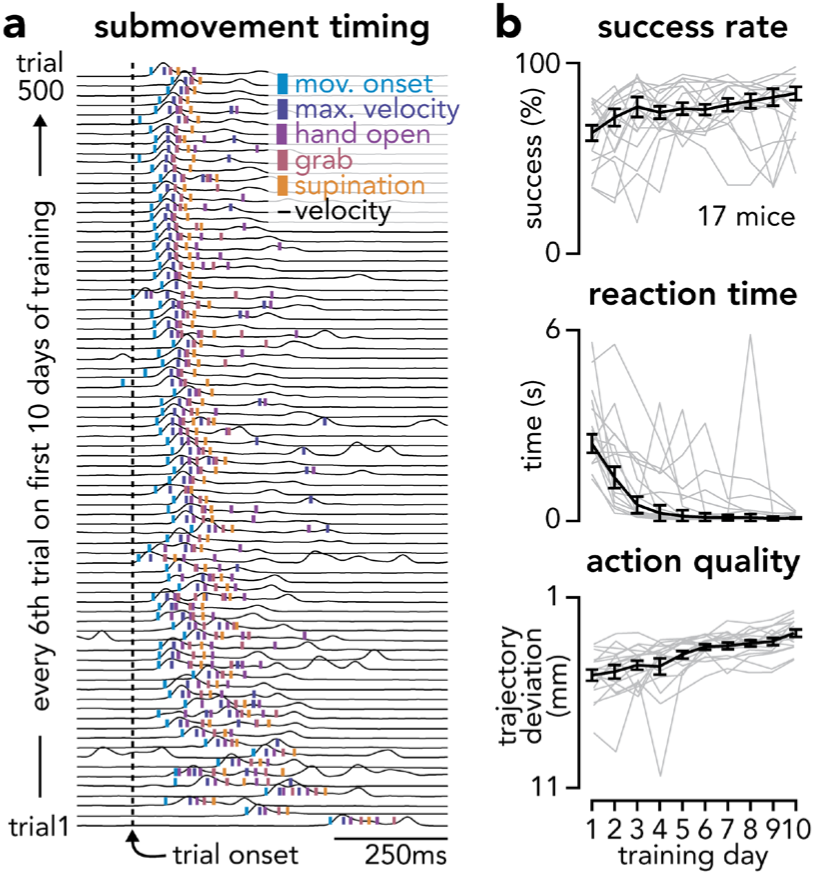
Skilled reach-to-grasp action learning. **(a)** Evolution of sub-movement timing during the first 10 days of training (50 trials / day, every 6th trial plotted) in an example animal, showing hand velocity overlaid with timing of movement onset, maximum velocity, hand open, grab, and supination aligned to trial onset. **(b)** Learning curves across 10 days of training (50 trials / day) for success rate (top), reaction time (middle), and action quality (bottom). Individual animals are shown in thin lines (*n* = 17 mice) with mean ± s.e.m. across animals overlaid in bold.

**Supplemental Figure 2:**
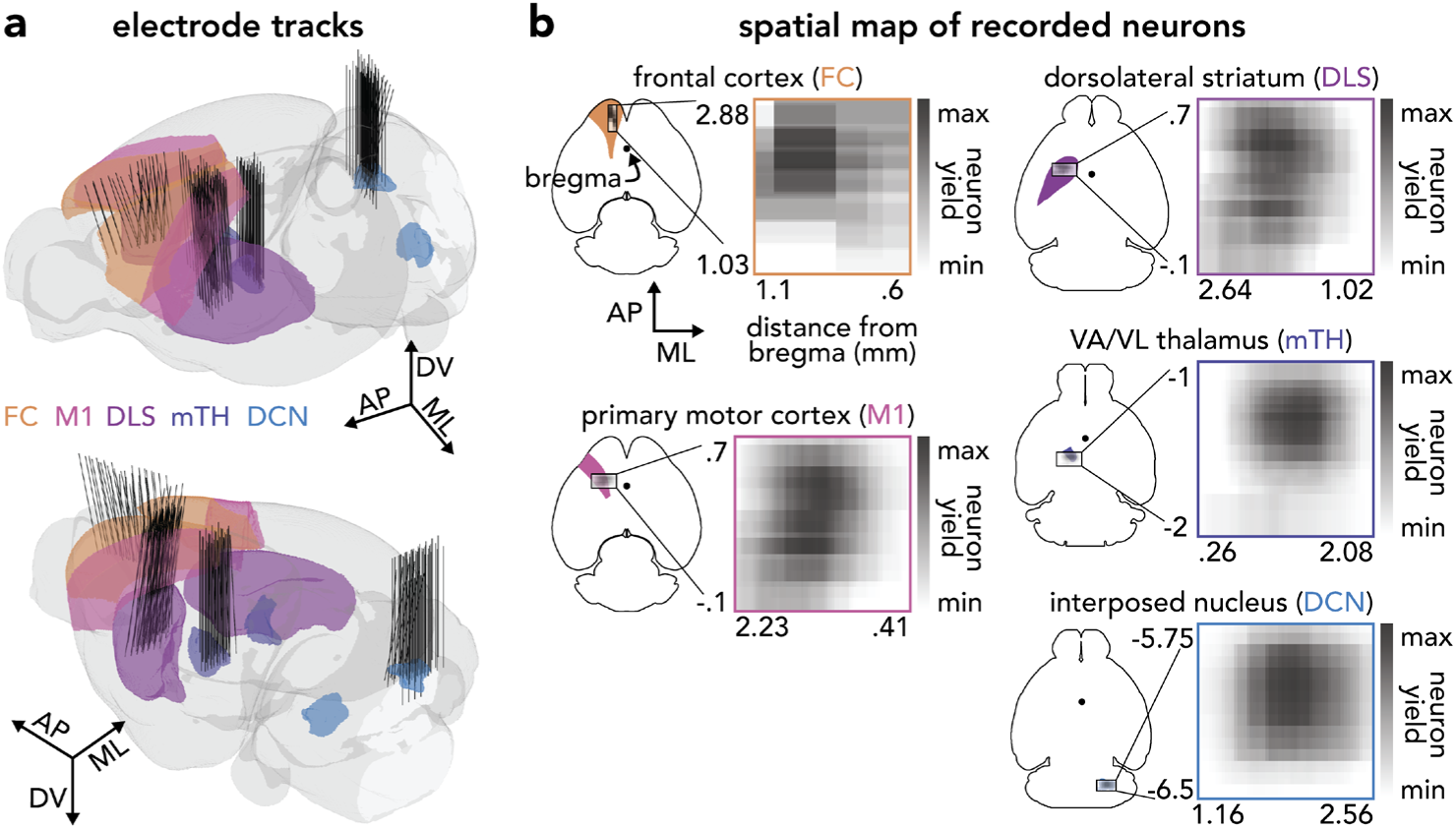
Anatomical localization of multi-area neural recordings. **(a)** Reconstruction of electrode shanks across recording sessions in 17 animals showing probe trajectories within each recorded brain area. **(b)** Spatial maps of single-neuron recording yield for each brain area across anterior-posterior and medial-lateral coordinates relative to bregma.

**Supplemental Figure 3:**
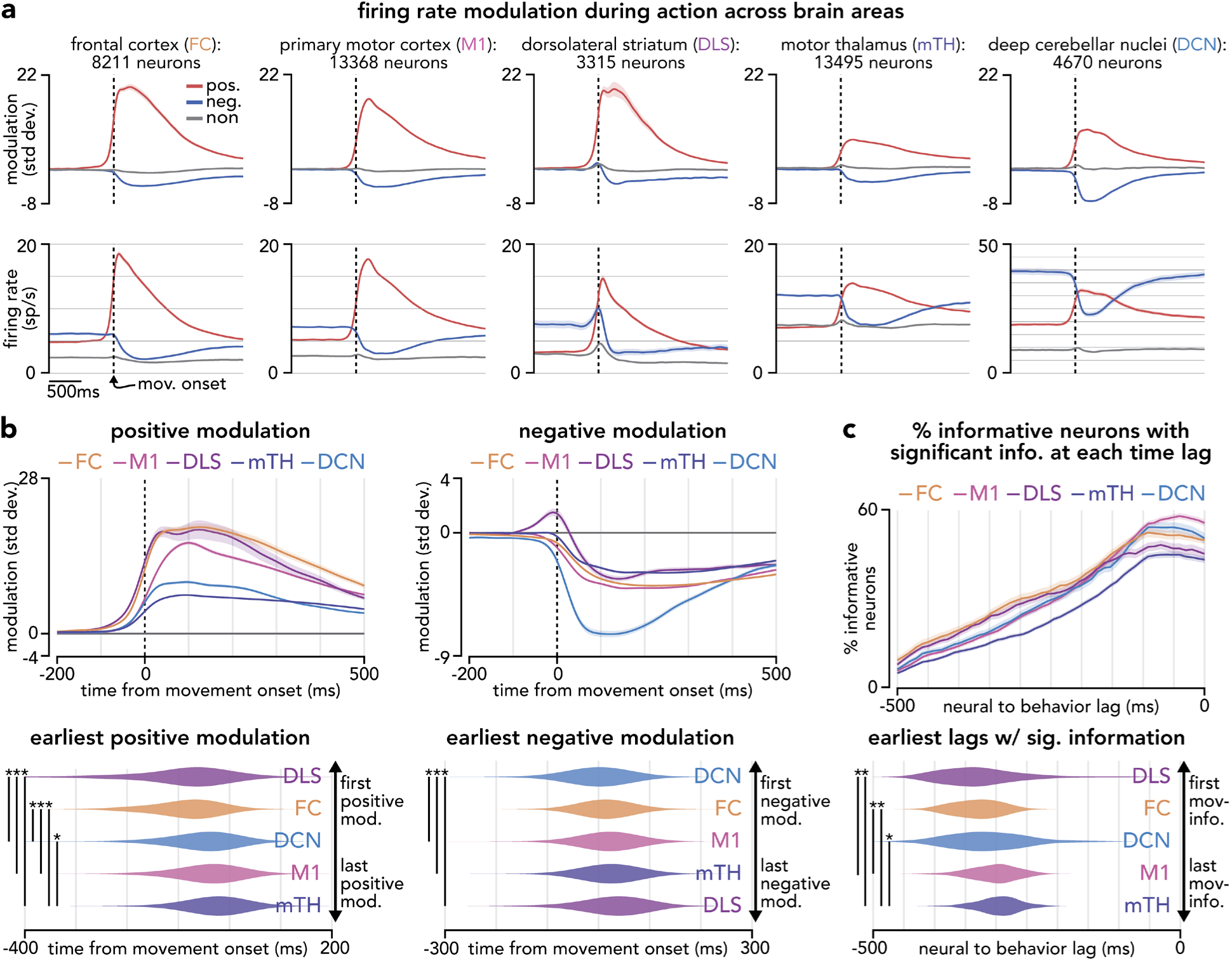
Cross-area firing rate modulation and action information timing. **(a)** Mean z-scored modulation (top) and firing rate (bottom) for positively modulated, negatively modulated, and non-modulated neurons during action production aligned to movement onset across recorded areas (line widths denote mean ± s.e.m. across sessions, FC: *n* = 54 recording sessions from 6 animals, M1: *n* = 138 sessions, 17 animals, DLS: *n* = 129 sessions, 17 animals, mTH: *n* = 95 sessions, 12 animals, DCN: *n* = 114 sessions, 16 animals). **(b)** Top: mean z-scored modulation in each area (line widths denote mean ± s.e.m. across sessions). Bottom: comparison of modulation onset timing across areas for positively and negatively modulated neurons (violin plots show session-level distributions, asterisks denote significant pairwise differences at *P*<0.05 with Bonferroni correction, two-sided Wilcoxon rank-sum test, session number in panel a). Positively modulated: DLS/FC: *P*=0.77, DLS/DCN: *P*<0.001, DLS/M1: *P*<0.001, DLS/mTH: *P*<0.001, FC/DCN: *P*<0.001, FC/M1: *P*<0.001, FC/mTH: *P*<0.001, DCN/M1: *P*=0.08, DCN/mTH: *P*<0.001, M1/mTH: *P*=0.02. Negatively modulated: DCN/FC: *P*=0.03, DCN/M1: *P*<0.001, DCN/mTH: *P*<0.001, DCN/DLS: *P*<0.001, FC/M1: *P*=0.11, FC/mTH: *P*=0.06, FC/DLS: *P*=0.01, M1/mTH: *P*=0.44, M1/DLS: *P*=0.05, mTH/DLS: *P*=0.13. **(c)** Top: percent of action-informative neurons with significant information at each neural-to-behavior time lag (line widths denote mean ± s.e.m. across sessions) and distributions of earliest significant information lags (bottom, violin plots show session-level distributions, asterisks denote significant pairwise differences at *P*<0.05 with Bonferroni correction, two-sided Wilcoxon rank-sum test, session number in panel a). DLS/FC: *P*=0.75, DLS/DCN: *P*=0.04, DLS/M1: *P*<0.001, DLS/mTH: *P*<0.001, FC/DCN: *P*=0.14, FC/M1: *P*<0.001, FC/mTH: *P*<0.001, DCN/M1: *P*=0.04, DCN/mTH: *P*=0.002, M1/mTH: *P*=0.11.

**Supplemental Figure 4:**
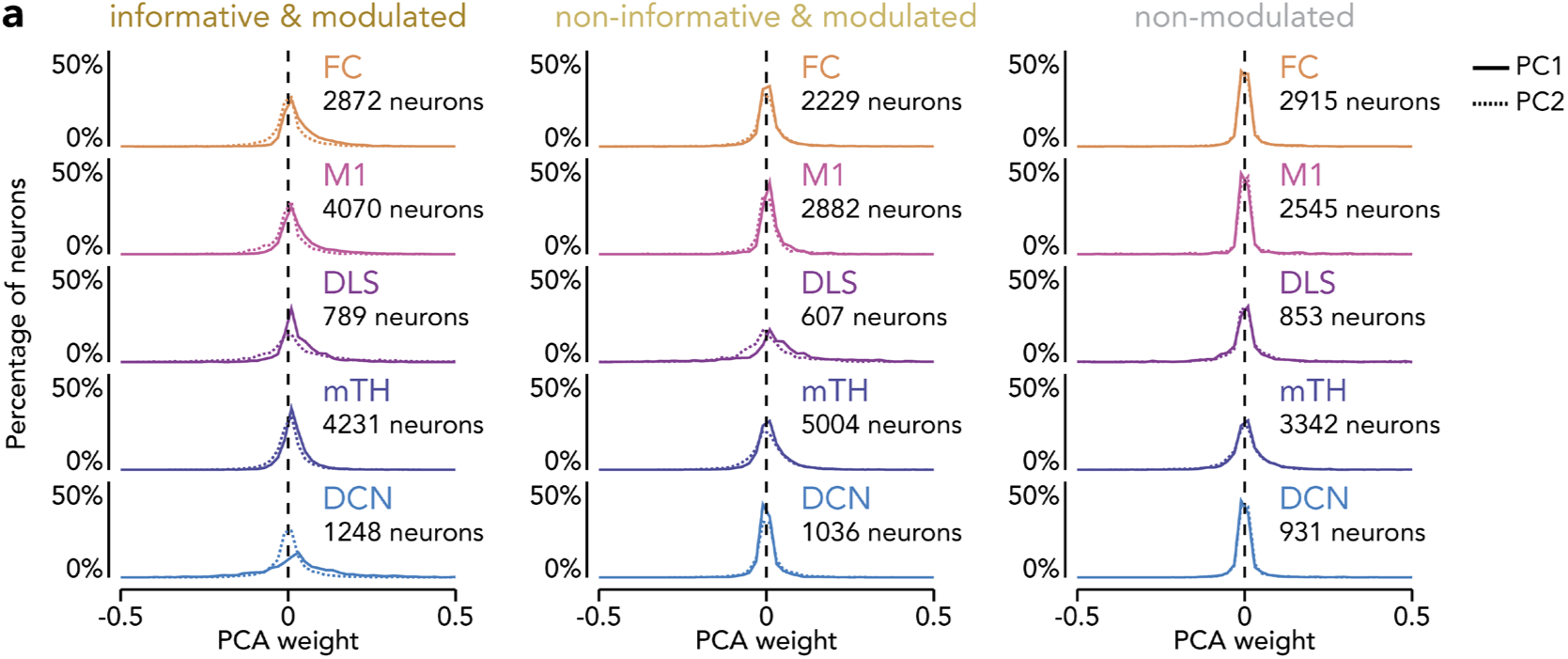
Distribution of PCA weights across brain areas. **(a)** Distribution of PCA weights (PC1 solid line, PC2 dashed line) across brain areas for PCAs computed from action-informative modulated neurons (*n* = 90 sessions across 13 animals meeting > 50 neuron criteria, including 13,210 neurons across areas), non-informative modulated neurons (*n* = 90 sessions, 13 animals, 11,758 neurons), and non-modulated neurons (*n* = 90 sessions, 13 animals, 10,586 neurons).

**Supplemental Figure 5:**
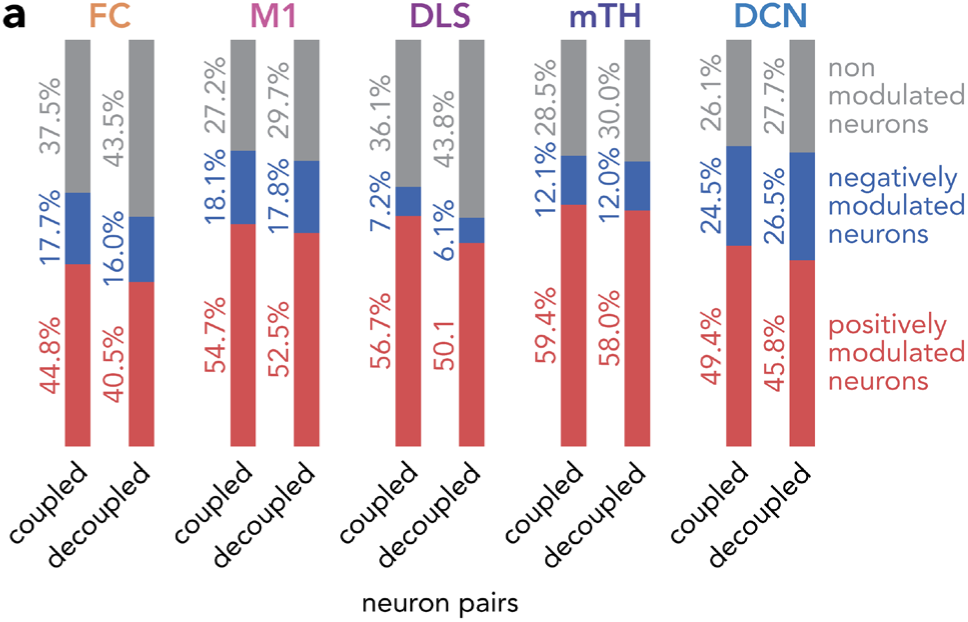
Action modulation within coupled and decoupled cross-area neuron pairs. **(a)** Proportion of positively modulated, negatively modulated, and non-modulated neurons within coupled and decoupled cross-area neuron pairs across brain areas.

**Supplemental Figure 6:**
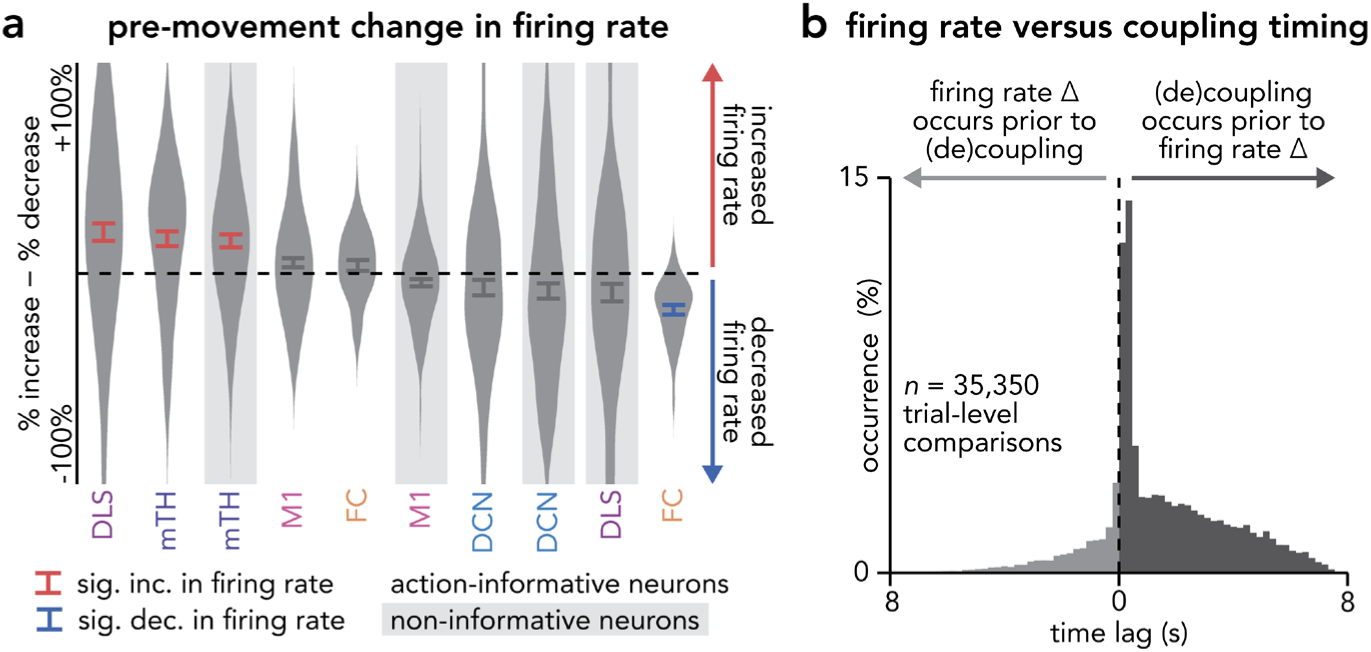
Pre-movement firing rate changes relative to network coupling and decoupling. **(a)** Difference between the percentage of neurons that increased versus decreased firing rate between early (−10 to −6.5 s) and late (−3.75 to −0.5 s) pre-movement periods for each neural population in each recorded area (violin plots show session-level distributions with mean ± s.e.m. overlaid). Red and blue bars indicate values significantly above or below zero and gray shaded background bars indicate distributions from non-informative neurons (*P*<0.05 with Bonferroni correction, one-sample *t*-test). Action-informative neurons: FC: *t*(38) = 1.50, *P*=0.14; M1: *t*(137) = 2.34, *P*=0.02; DLS: *t*(128) = 4.71, *P*<0.001; mTH: *t*(94) = 4.73, *P*<0.001; DCN: *t*(113) = −1.81, *P*=0.07. Non-informative neurons: FC: *t*(38) = −7.65, *P*<0.001; M1: *t*(137) = −2.53, *P*=0.01; DLS: *t*(128) = −2.29, *P*=0.02; mTH: *t*(94) = 5.07, *P*<0.001; DCN: *t*(113) = −2.17, *P*=0.03. FC: *n* = 55 recording sessions from 6 animals; M1: *n* = 138 sessions, 17 animals; DLS: *n* = 129 sessions, 17 animals; mTH: *n* = 95 sessions, 12 animals; DCN: *n* = 114 sessions, 16 animals. **(b)** Distribution of time lags between the onset of pre-movement firing rate changes and the onset of coupling or decoupling across neuron pairs on each trial, pooled across sessions and area pairs.

**Supplemental Figure 7:**
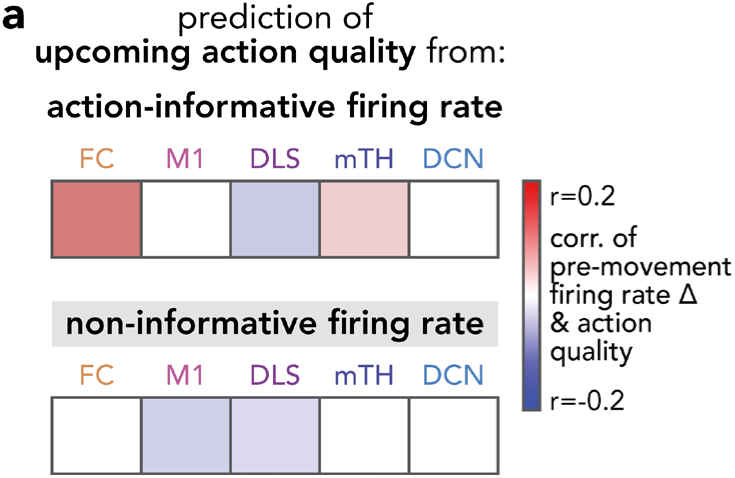
Relationship between pre-movement firing rate changes and action quality. **(a)** Trial-by-trial correlations between action quality and pre-movement changes in firing rate of action-informative neurons (top) and non-informative neurons (bottom). Color indicates significant correlation at *P*<0.05, Bonferroni-corrected (Spearman correlation). Action-informative neurons: FC: *r* = 0.13, *P*<0.001; M1: *r* = −0.02, *P*=0.04; DLS: *r* = −0.05, *P*<0.001; mTH: *r* = 0.05, *P*<0.001; DCN: *r* = −0.03, *P*=0.01. Non-informative neurons: FC: *r* = −0.01, *P*=0.42; M1: *r* = −0.05, *P*<0.001; DLS: *r* = −0.04, *P*=0.001; mTH: *r* = 0.02, *P*=0.12; DCN: *r* = −0.02, *P*=0.12. FC: *n* = 3768 trials from 6 animals; M1: *n* = 9380 trials from 17 animals; DLS: *n* = 8752 trials from 17 animals; mTH: *n* = 6782 trials from 12 animals; DCN: *n* = 7733 trials from 16 animals.

**Supplemental Figure 8:**
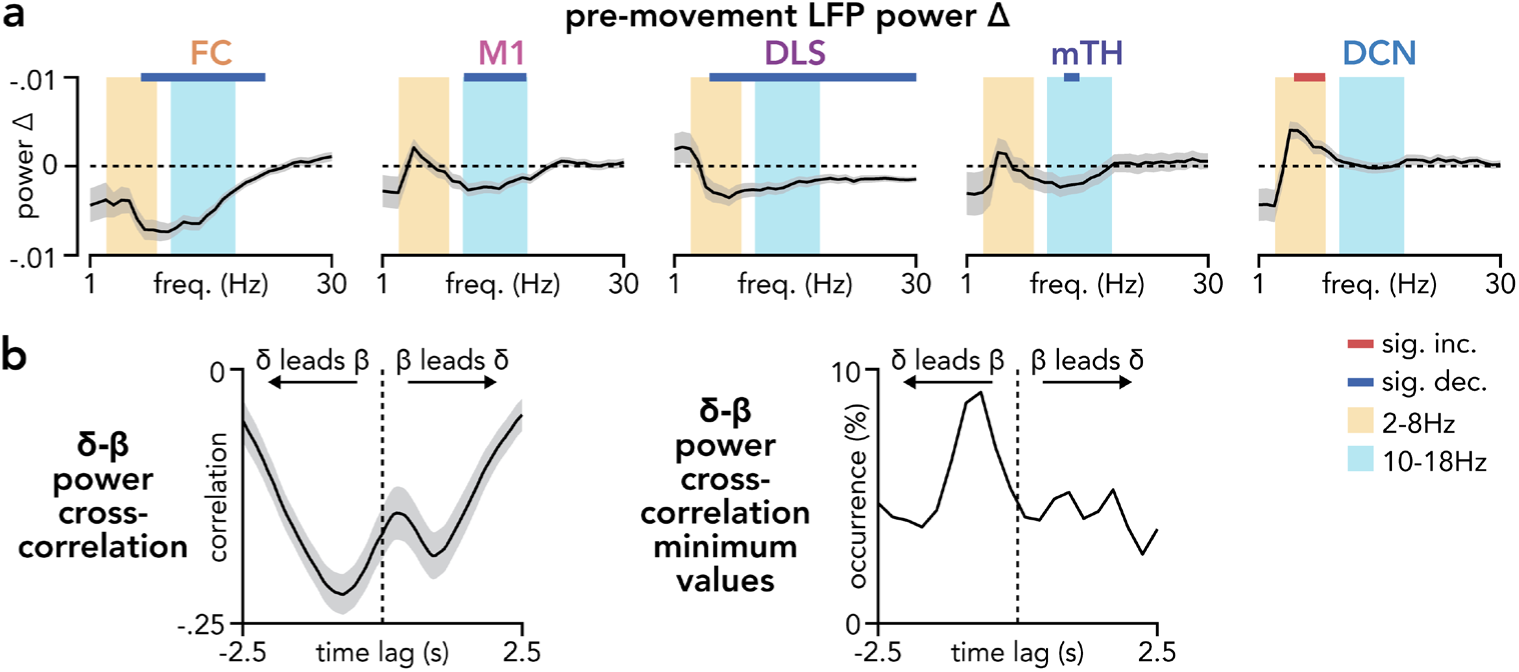
Pre-movement LFP power changes across brain areas. **(a)** Changes in pre-movement LFP power across all recorded areas. Power spectra show significant increases (red bars) or decreases (blue bars) between early and late pre-movement periods, highlighting 2-8 Hz (yellow background) and 10-18 Hz (blue background) frequency bands (two-sided Wilcoxon signed-rank test, Bonferroni-corrected, P < 0.05). Line width denotes mean ± s.e.m. across sessions (FC: *n* = 55 sessions across 6 animals; M1: *n* = 155 sessions across 17 animals; DLS: *n* = 155 sessions across 17 animals; mTH: *n* = 100 sessions across 12 animals; DCN: *n* = 145 sessions across 16 animals). **(b)** δ-β power coordination during the pre-movement period. Left: average cross-correlation between δ power in DCN and β power in each other area (M1, DLS, mTH, FC), for sessions with pre-movement δ power increases and β power decreases, computed separately for each area within each animal (line width represents mean ± s.e.m. across areas pooled across animals, n = 49 cross-correlations pooled across 16 animals). Right: distribution of lag values at which each cross-correlation reached its minimum (strongest anti-correlation).

**Supplemental Figure 9:**
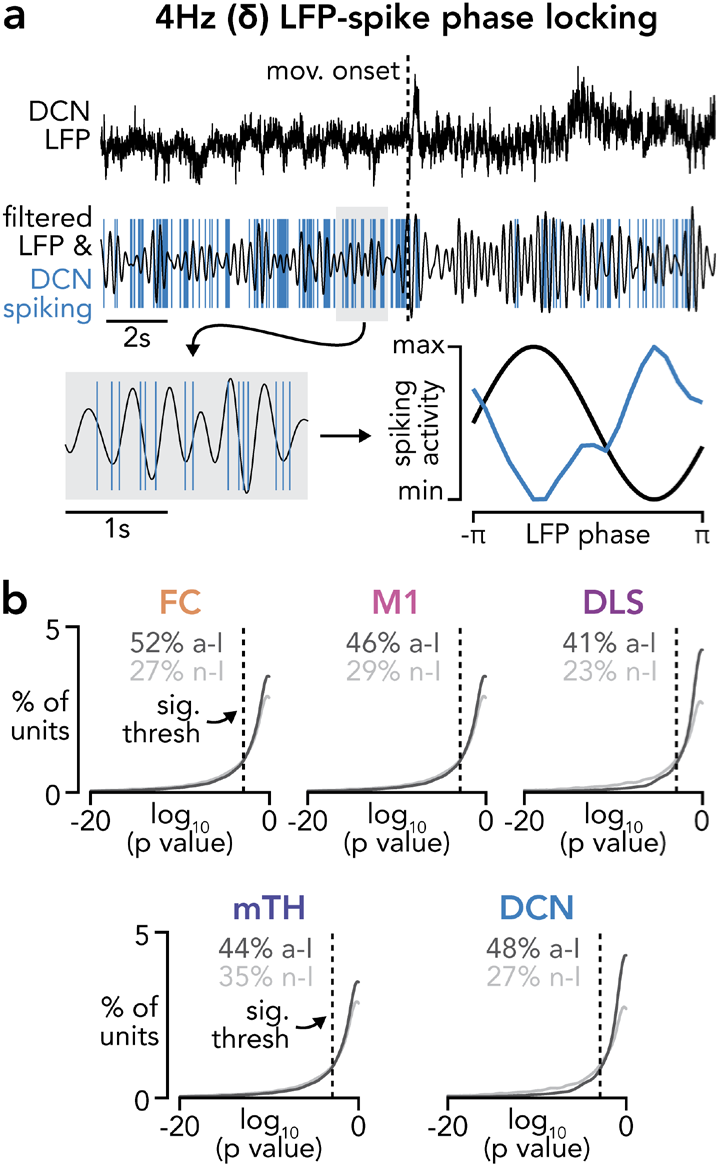
Pre-movement δ-band LFP-spike phase locking across brain areas. **(a)** Example raw LFP from DCN during a single reaching trial and corresponding δ-band (4 Hz) filtered LFP with overlaid spike times from an example DCN neuron. Corresponding polar histogram showing the distribution of spike phases for the same neuron during the pre-movement period. **(b)** Percentage of units significantly phase-locked to δ-band LFP across areas. Plots show cumulative distributions of log-transformed Rayleigh test *P* values for all recorded units. Dashed lines indicate significance threshold (*P* < 0.05). Summary statistics above each plot report the percentage of significantly phase-locked action-informative (a-I) and non-informative (n-I) units (FC: *n* = 28,728 action-informative / 36,960 non-informative neuron-LFP pairs from 54 sessions across 6 animals; M1: *n* = 26,688 / 26,784 neuron-LFP pairs, 138 sessions, 17 animals; DLS: *n* = 5,968 / 7,292 neuron-LFP pairs, 129 sessions, 17 animals; mTH: *n* = 25,576 / 37,424 neuron-LFP pairs, 97 sessions, 12 animals; DCN: *n* = 12,508 / 15,948 neuron-LFP pairs, 114 sessions, 16 animals).

**Supplemental Figure 10:**
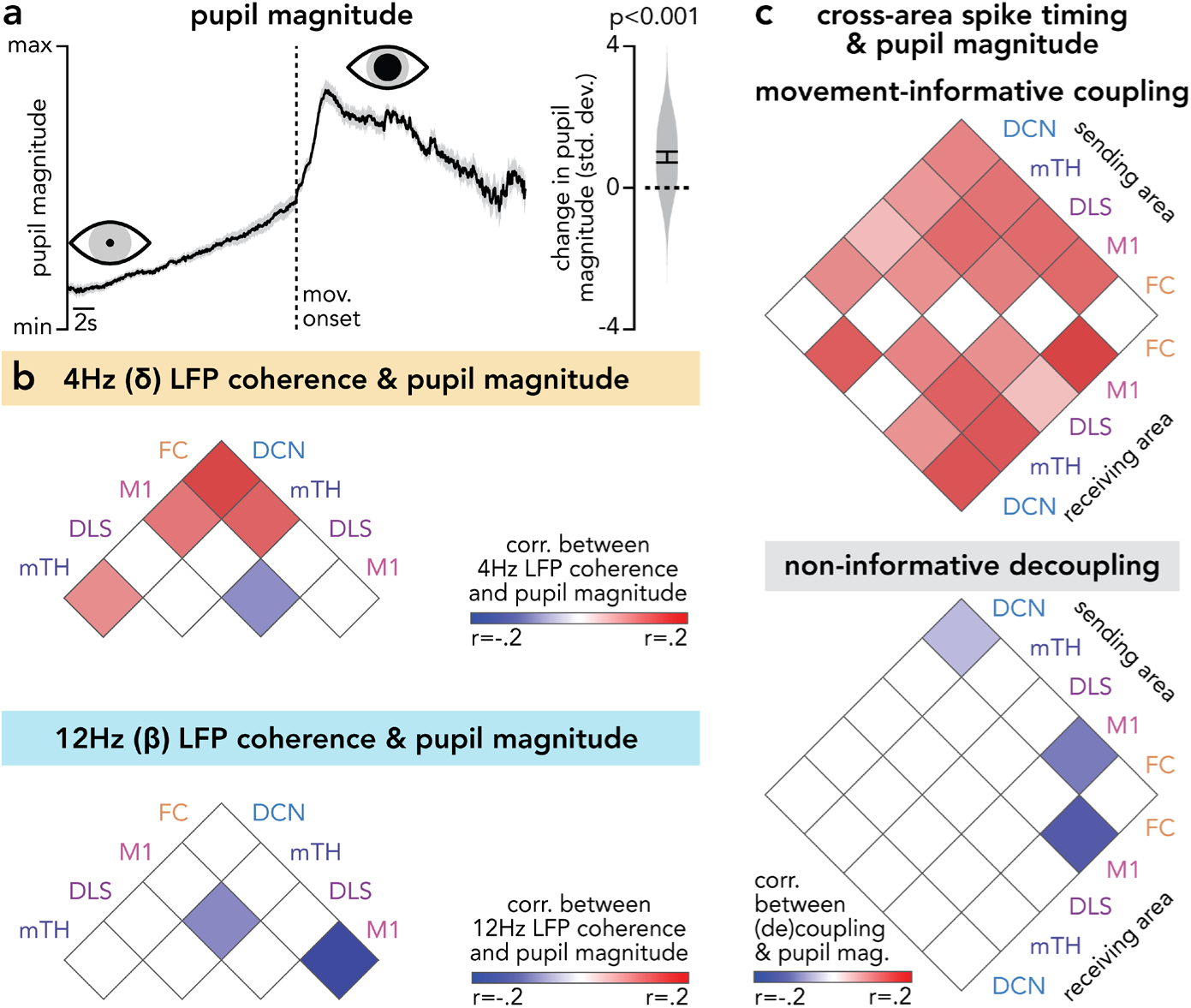
Pupil dynamics are correlated with coupling/decoupling and LFP coherence. **(a)** Average pupil magnitude aligned to movement onset (left) and summary of pre-movement pupil magnitude changes across sessions (right). Pupil magnitude increased from early to late pre-movement periods (*t*(29) = 5.60, *P*<0.001, one-sample t-test, *n* = 30 sessions from 5 animals). **(b)** Matrices of trial-by-trial correlations between pupil magnitude and pre-movement LFP coherence at 4 Hz (δ, top) and 12 Hz (β, bottom) across area pairs. Color indicates significance at *P*<0.05, Bonferroni-corrected (Spearman correlation). Detailed statistics in Supp. Table 4. **(c)** Matrices of trial-by-trial correlations between pupil magnitude and action-informative coupling (top) or non-informative decoupling (bottom) for all directional area pairs. Color indicates significance at *P*<0.05, Bonferroni-corrected (Spearman correlation). Detailed statistics in Supp. Table 4.

**Supplemental Table 1.** Cross-area coupling and decoupling statistics.

| area pair sorted by coupling | *P* value | *t* statistic | *n* sessions |
| --- | --- | --- | --- |
| **action-informative neuron pairs** |  |  |  |
| DCN → DLS | < 0.001 | *t*(108) = 7.52 | 109 |
| M1 → DLS | < 0.001 | *t*(125) = 8.44 | 126 |
| DCN → mTH | < 0.001 | *t*(83) = 8.85 | 84 |
| DCN → FC | < 0.001 | *t*(45) = 5.44 | 46 |
| DLS → FC | < 0.001 | *t*(50) = 5.01 | 51 |
| DCN → M1 | < 0.001 | *t*(112) = 7.58 | 113 |
| DLS → mTH | < 0.001 | *t*(91) = 5.50 | 92 |
| M1 → mTH | < 0.001 | *t*(92) = 8.28 | 93 |
| M1 → FC | < 0.001 | *t*(53) = 6.69 | 54 |
| FC → DLS | < 0.001 | *t*(50) = 4.20 | 51 |
| DLS → DCN | < 0.001 | *t*(108) = 5.56 | 109 |
| M1 → DCN | < 0.001 | *t*(112) = 6.53 | 113 |
| DLS → M1 | < 0.001 | *t*(125) = 4.79 | 126 |
| mTH → FC | < 0.001 | *t*(50) = 5.48 | 51 |
| mTH → DLS | < 0.001 | *t*(91) = 3.84 | 92 |
| FC → mTH | 0.010 | *t*(50) = 2.68 | 51 |
| mTH → M1 | 0.011 | *t*(92) = 2.61 | 93 |
| mTH → DCN | 0.077 | *t*(83) = 1.79 | 84 |
| FC → M1 | 0.258 | *t*(53) = 1.14 | 54 |
| FC → DCN | 0.272 | *t*(45) = -1.11 | 46 |
| **non-informative neuron pairs** |  |  |  |
| DCN → mTH | < 0.001 | *t*(83) = 8.22 | 84 |
| DCN → FC | < 0.001 | *t*(45) = 4.32 | 46 |
| M1 → DCN | < 0.001 | *t*(112) = 5.76 | 113 |
| M1 → mTH | < 0.001 | *t*(92) = 4.83 | 93 |
| DCN → M1 | < 0.001 | *t*(112) = 3.56 | 113 |
| DLS → DCN | 0.022 | *t*(108) = 2.33 | 109 |
| DCN → DLS | 0.414 | *t*(108) = 0.82 | 109 |
| M1 → DLS | 0.444 | *t*(126) = 0.77 | 127 |
| mTH → FC | 0.495 | *t*(50) = 0.69 | 51 |
| mTH → DCN | 0.566 | *t*(83) = -0.58 | 84 |
| M1 → FC | 0.652 | *t*(53) = -0.45 | 54 |
| mTH → M1 | 0.105 | *t*(92) = -1.64 | 93 |
| mTH → DLS | 0.001 | *t*(91) = -3.29 | 92 |
| DLS → mTH | 0.008 | *t*(91) = -2.72 | 92 |
| DLS → M1 | < 0.001 | *t*(126) = -3.95 | 127 |
| DLS → FC | < 0.001 | *t*(50) = -4.34 | 51 |
| FC → mTH | < 0.001 | *t*(50) = -5.28 | 51 |
| FC → DCN | < 0.001 | *t*(45) = -6.10 | 46 |
| FC → M1 | < 0.001 | *t*(53) = -6.92 | 54 |
| FC → DLS | < 0.001 | *t*(50) = -6.11 | 51 |

**Supplemental Table 2.** Coupling and decoupling correlations to action quality and neural dynamics.

| Prediction of upcoming action quality from: |  |  |  |
| --- | --- | --- | --- |
| area pair | action-informative coupling | non-informative decoupling | *n* |
| DCN → FC | $r = -0.07, P < 0.001$ | $r = 0.04, P = 0.037$ | 2379 trials, 46 sessions, 6 animals |
| DCN → M1 | $r = -0.09, P < 0.001$ | $r = 0.02, P = 0.106$ | 5154 trials, 104 sessions, 15 animals |
| DCN → DLS | $r = -0.07, P < 0.001$ | $r = 0.05, P < 0.001$ | 5006 trials, 100 sessions, 15 animals |
| DCN → mTH | $r = -0.10, P < 0.001$ | $r = 0.01, P = 0.429$ | 4164 trials, 84 sessions, 12 animals |
| mTH → FC | $r = -0.14, P < 0.001$ | $r = 0.08, P < 0.001$ | 2752 trials, 53 sessions, 6 animals |
| mTH → M1 | $r = -0.15, P < 0.001$ | $r = 0.02, P = 0.117$ | 4690 trials, 95 sessions, 12 animals |
| mTH → DLS | $r = -0.13, P < 0.001$ | $r = 0.04, P = 0.004$ | 4580 trials, 92 sessions, 12 animals |
| mTH → DCN | $r = -0.07, P < 0.001$ | $r = -0.01, P = 0.553$ | 4164 trials, 84 sessions, 12 animals |
| DLS → FC | $r = -0.15, P < 0.001$ | $r = 0.07, P < 0.001$ | 2675 trials, 51 sessions, 6 animals |
| DLS → M1 | $r = -0.10, P < 0.001$ | $r = 0.07, P < 0.001$ | 5693 trials, 115 sessions, 16 animals |
| DLS → mTH | $r = -0.15, P < 0.001$ | $r = 0.05, P < 0.001$ | 4580 trials, 92 sessions, 12 animals |
| DLS → DCN | $r = -0.06, P < 0.001$ | $r = 0.02, P = 0.155$ | 5006 trials, 100 sessions, 15 animals |
| M1 → FC | $r = -0.19, P < 0.001$ | $r = 0.17, P < 0.001$ | 2814 trials, 54 sessions, 6 animals |
| M1 → DLS | $r = -0.11, P < 0.001$ | $r = 0.08, P < 0.001$ | 5693 trials, 115 sessions, 16 animals |
| M1 → mTH | $r = -0.16, P < 0.001$ | $r = 0.02, P = 0.125$ | 4690 trials, 95 sessions, 12 animals |
| M1 → DCN | $r = -0.05, P < 0.001$ | $r = 0.00, P = 0.951$ | 5154 trials, 104 sessions, 15 animals |
| FC → M1 | $r = -0.22, P < 0.001$ | $r = 0.23, P < 0.001$ | 2814 trials, 54 sessions, 6 animals |
| FC → DLS | $r = -0.14, P < 0.001$ | $r = 0.11, P < 0.001$ | 2675 trials, 51 sessions, 6 animals |
| FC → mTH | $r = -0.16, P < 0.001$ | $r = 0.14, P < 0.001$ | 2752 trials, 53 sessions, 6 animals |
| FC → DCN | $r = -0.06, P = 0.003$ | $r = 0.04, P = 0.043$ | 2379 trials, 46 sessions, 6 animals |
| Prediction of upcoming neural trajectory consistency from: |  |  |  |
| area pair | action-informative coupling | non-informative decoupling | *n* |
| DCN → FC | $r = 0.09, P < 0.001$ | $r = -0.00, P = 0.970$ | 2379 trials, 46 sessions, 6 animals |
| DCN → M1 | $r = 0.10, P < 0.001$ | $r = 0.01, P = 0.687$ | 5154 trials, 104 sessions, 15 animals |
| DCN → DLS | $r = 0.07, P < 0.001$ | $r = 0.00, P = 0.912$ | 5006 trials, 100 sessions, 15 animals |
| DCN → mTH | $r = 0.10, P < 0.001$ | $r = 0.02, P = 0.112$ | 4164 trials, 84 sessions, 12 animals |
| mTH → FC | $r = 0.10, P < 0.001$ | $r = -0.09, P < 0.001$ | 2752 trials, 53 sessions, 6 animals |
| mTH → M1 | $r = 0.09, P < 0.001$ | $r = -0.02, P = 0.106$ | 4690 trials, 95 sessions, 12 animals |
| mTH → DLS | $r = 0.10, P < 0.001$ | $r = -0.02, P = 0.097$ | 4580 trials, 92 sessions, 12 animals |
| mTH → DCN | $r = 0.07, P < 0.001$ | $r = 0.03, P = 0.105$ | 4164 trials, 84 sessions, 12 animals |
| DLS → FC | $r = 0.14, P < 0.001$ | $r = -0.06, P = 0.003$ | 2675 trials, 51 sessions, 6 animals |
| DLS → M1 | $r = 0.08, P < 0.001$ | $r = -0.05, P < 0.001$ | 5693 trials, 115 sessions, 16 animals |
| DLS → mTH | $r = 0.13, P < 0.001$ | $r = -0.02, P = 0.128$ | 4580 trials, 92 sessions, 12 animals |
| DLS → DCN | $r = 0.06, P < 0.001$ | $r = 0.03, P = 0.069$ | 5006 trials, 100 sessions, 15 animals |
| M1 → FC | $r = 0.15, P < 0.001$ | $r = -0.15, P < 0.001$ | 2814 trials, 54 sessions, 6 animals |
| M1 → DLS | $r = 0.08, P < 0.001$ | $r = -0.04, P = 0.001$ | 5693 trials, 115 sessions, 16 animals |
| M1 → mTH | $r = 0.12, P < 0.001$ | $r = -0.02, P = 0.285$ | 4690 trials, 95 sessions, 12 animals |
| M1 → DCN | $r = 0.06, P < 0.001$ | $r = 0.03, P = 0.044$ | 5154 trials, 104 sessions, 15 animals |
| FC → M1 | $r = 0.15, P < 0.001$ | $r = -0.22, P < 0.001$ | 2814 trials, 54 sessions, 6 animals |
| FC → DLS | $r = 0.13, P < 0.001$ | $r = -0.11, P < 0.001$ | 2675 trials, 51 sessions, 6 animals |
| FC → mTH | $r = 0.15, P < 0.001$ | $r = -0.16, P < 0.001$ | 2752 trials, 53 sessions, 6 animals |
| FC → DCN | $r = 0.07, P < 0.001$ | $r = -0.02, P = 0.294$ | 2379 trials, 46 sessions, 6 animals |

**Supplemental Table 3.**
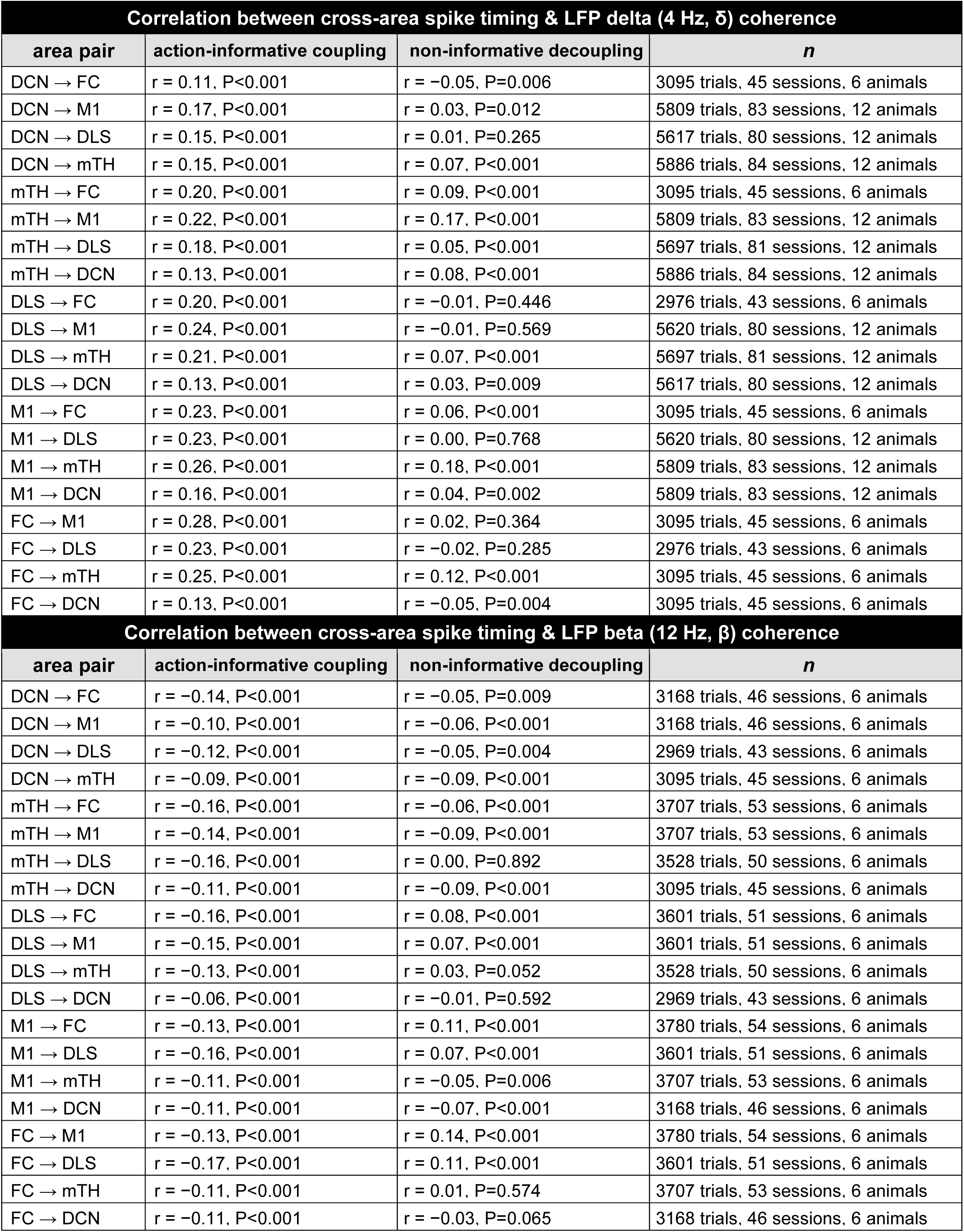
LFP coherence correlations to coupling and decoupling.

| Correlation between cross-area spike timing & LFP delta (4 Hz, $\delta$) coherence | | | |
| --- | --- | --- | --- |
| area pair | action-informative coupling | non-informative decoupling | *n* |
| DCN → FC | $r = 0.11, P < 0.001$ | $r = -0.05, P = 0.006$ | 3095 trials, 45 sessions, 6 animals |
| DCN → M1 | $r = 0.17, P < 0.001$ | $r = 0.03, P = 0.012$ | 5809 trials, 83 sessions, 12 animals |
| DCN → DLS | $r = 0.15, P < 0.001$ | $r = 0.01, P = 0.265$ | 5617 trials, 80 sessions, 12 animals |
| DCN → mTH | $r = 0.15, P < 0.001$ | $r = 0.07, P < 0.001$ | 5886 trials, 84 sessions, 12 animals |
| mTH → FC | $r = 0.20, P < 0.001$ | $r = 0.09, P < 0.001$ | 3095 trials, 45 sessions, 6 animals |
| mTH → M1 | $r = 0.22, P < 0.001$ | $r = 0.17, P < 0.001$ | 5809 trials, 83 sessions, 12 animals |
| mTH → DLS | $r = 0.18, P < 0.001$ | $r = 0.05, P < 0.001$ | 5697 trials, 81 sessions, 12 animals |
| mTH → DCN | $r = 0.13, P < 0.001$ | $r = 0.08, P < 0.001$ | 5886 trials, 84 sessions, 12 animals |
| DLS → FC | $r = 0.20, P < 0.001$ | $r = -0.01, P = 0.446$ | 2976 trials, 43 sessions, 6 animals |
| DLS → M1 | $r = 0.24, P < 0.001$ | $r = -0.01, P = 0.569$ | 5620 trials, 80 sessions, 12 animals |
| DLS → mTH | $r = 0.21, P < 0.001$ | $r = 0.07, P < 0.001$ | 5697 trials, 81 sessions, 12 animals |
| DLS → DCN | $r = 0.13, P < 0.001$ | $r = 0.03, P = 0.009$ | 5617 trials, 80 sessions, 12 animals |
| M1 → FC | $r = 0.23, P < 0.001$ | $r = 0.06, P < 0.001$ | 3095 trials, 45 sessions, 6 animals |
| M1 → DLS | $r = 0.23, P < 0.001$ | $r = 0.00, P = 0.768$ | 5620 trials, 80 sessions, 12 animals |
| M1 → mTH | $r = 0.26, P < 0.001$ | $r = 0.18, P < 0.001$ | 5809 trials, 83 sessions, 12 animals |
| M1 → DCN | $r = 0.16, P < 0.001$ | $r = 0.04, P = 0.002$ | 5809 trials, 83 sessions, 12 animals |
| FC → M1 | $r = 0.28, P < 0.001$ | $r = 0.02, P = 0.364$ | 3095 trials, 45 sessions, 6 animals |
| FC → DLS | $r = 0.23, P < 0.001$ | $r = -0.02, P = 0.285$ | 2976 trials, 43 sessions, 6 animals |
| FC → mTH | $r = 0.25, P < 0.001$ | $r = 0.12, P < 0.001$ | 3095 trials, 45 sessions, 6 animals |
| FC → DCN | $r = 0.13, P < 0.001$ | $r = -0.05, P = 0.004$ | 3095 trials, 45 sessions, 6 animals |
| Correlation between cross-area spike timing & LFP beta (12 Hz, $\beta$) coherence | | | |
| area pair | action-informative coupling | non-informative decoupling | *n* |
| DCN → FC | $r = -0.14, P < 0.001$ | $r = -0.05, P = 0.009$ | 3168 trials, 46 sessions, 6 animals |
| DCN → M1 | $r = -0.10, P < 0.001$ | $r = -0.06, P < 0.001$ | 3168 trials, 46 sessions, 6 animals |
| DCN → DLS | $r = -0.12, P < 0.001$ | $r = -0.05, P = 0.004$ | 2969 trials, 43 sessions, 6 animals |
| DCN → mTH | $r = -0.09, P < 0.001$ | $r = -0.09, P < 0.001$ | 3095 trials, 45 sessions, 6 animals |
| mTH → FC | $r = -0.16, P < 0.001$ | $r = -0.06, P < 0.001$ | 3707 trials, 53 sessions, 6 animals |
| mTH → M1 | $r = -0.14, P < 0.001$ | $r = -0.09, P < 0.001$ | 3707 trials, 53 sessions, 6 animals |
| mTH → DLS | $r = -0.16, P < 0.001$ | $r = 0.00, P = 0.892$ | 3528 trials, 50 sessions, 6 animals |
| mTH → DCN | $r = -0.11, P < 0.001$ | $r = -0.09, P < 0.001$ | 3095 trials, 45 sessions, 6 animals |
| DLS → FC | $r = -0.16, P < 0.001$ | $r = 0.08, P < 0.001$ | 3601 trials, 51 sessions, 6 animals |
| DLS → M1 | $r = -0.15, P < 0.001$ | $r = 0.07, P < 0.001$ | 3601 trials, 51 sessions, 6 animals |
| DLS → mTH | $r = -0.13, P < 0.001$ | $r = 0.03, P = 0.052$ | 3528 trials, 50 sessions, 6 animals |
| DLS → DCN | $r = -0.06, P < 0.001$ | $r = -0.01, P = 0.592$ | 2969 trials, 43 sessions, 6 animals |
| M1 → FC | $r = -0.13, P < 0.001$ | $r = 0.11, P < 0.001$ | 3780 trials, 54 sessions, 6 animals |
| M1 → DLS | $r = -0.16, P < 0.001$ | $r = 0.07, P < 0.001$ | 3601 trials, 51 sessions, 6 animals |
| M1 → mTH | $r = -0.11, P < 0.001$ | $r = -0.05, P = 0.006$ | 3707 trials, 53 sessions, 6 animals |
| M1 → DCN | $r = -0.11, P < 0.001$ | $r = -0.07, P < 0.001$ | 3168 trials, 46 sessions, 6 animals |
| FC → M1 | $r = -0.13, P < 0.001$ | $r = 0.14, P < 0.001$ | 3780 trials, 54 sessions, 6 animals |
| FC → DLS | $r = -0.17, P < 0.001$ | $r = 0.11, P < 0.001$ | 3601 trials, 51 sessions, 6 animals |
| FC → mTH | $r = -0.11, P < 0.001$ | $r = 0.01, P = 0.574$ | 3707 trials, 53 sessions, 6 animals |
| FC → DCN | $r = -0.11, P < 0.001$ | $r = -0.03, P = 0.065$ | 3168 trials, 46 sessions, 6 animals |

**Supplemental Table 4.**
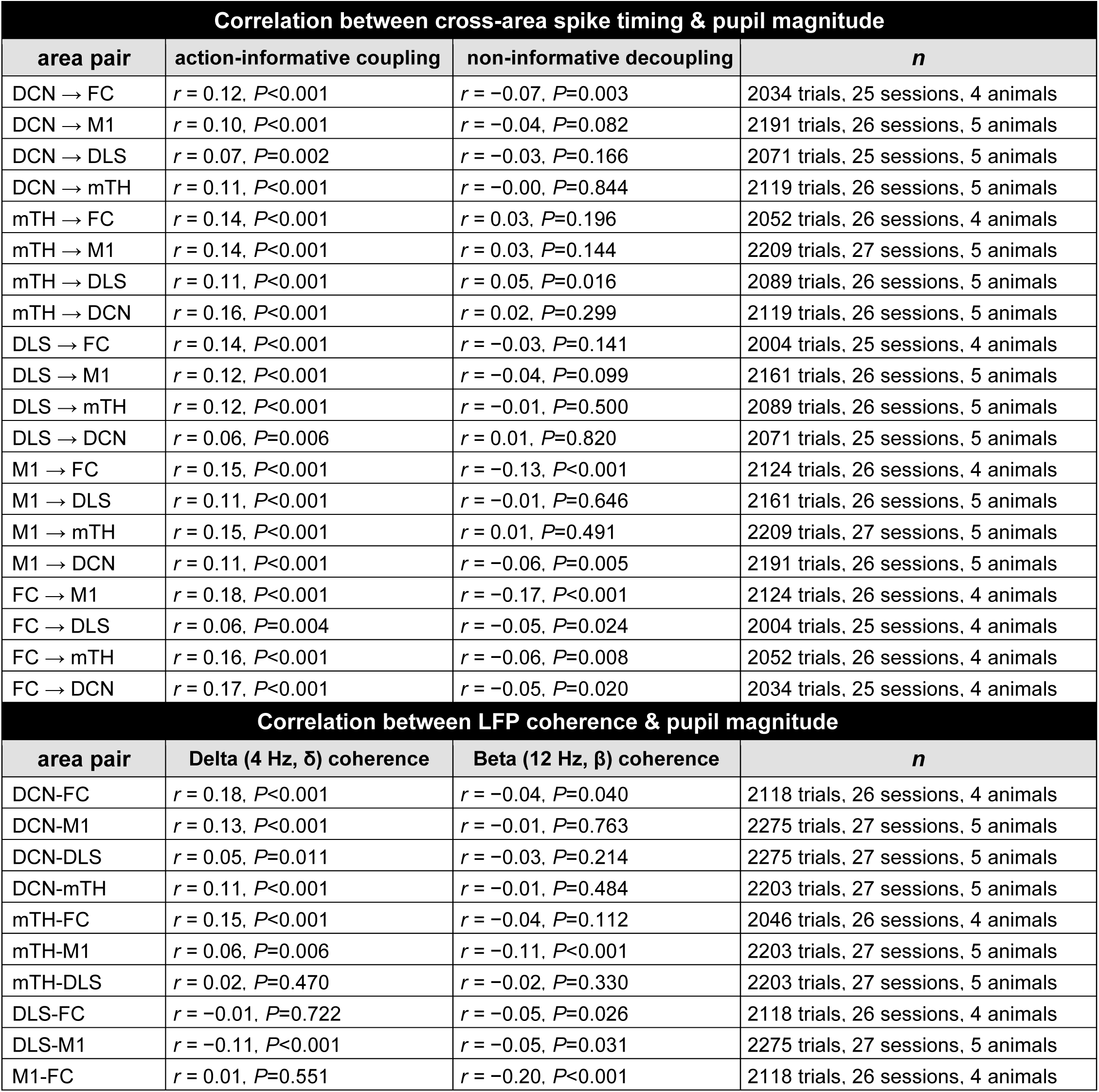
Pupil magnitude correlations to coupling and decoupling and LFP coherence.

## Methods

### Animals

All experimental procedures were approved by the Institutional Animal Care and Use Committee at the University of North Carolina at Chapel Hill. Wild-type and *Slc32a1-ChR2(H134R)-*EYFP mice (The Jackson Laboratory Strain #:014548) of both sexes were studied. A total of 23 mice were used (17 wild-type and 6 *Slc32a1-ChR2(H134R)-*EYFP).

### Reach-to-grasp task

Adult mice (12-25 week old) naive to any behavioral task were surgically implanted with 3D-printed headposts (Formlabs; adapted from the RIVETS headpost^50^) directly on the skull. Briefly, mice were anesthetized with isoflurane and placed in a stereotactic frame (Kopf Instruments) on a 37°C heating pad. Aseptic technique was used during surgical procedures. The scalp and periosteum over the skull were removed, a layer of UV-curing OptiBond adhesive (Kerr Dental) was applied, and a headpost was affixed with Calibra Universal Self-Adhesive Resin Cement (Dentsply Sirona). A thin layer of dental cement was placed over the exposed skull at the center of the headpost. After a minimum of 10 days of recovery, mice were food-restricted to 75-85% baseline weight and trained on a head-fixed single-pellet reach-to-grasp task. In the reach-to-grasp task, mice reach out and grab a food pellet (20mg, Dustless Precision Pellets, Purified; Bio-Serv). Initial training on this task began with mice licking pellets placed directly below their mouth, often using their hand to guide the pellet into their mouth. Pellets arrived ∼200 ms after the start of an auditory tone (5 kHz) by rotating a turntable driven by custom-programmed Arduino software, as previously used^15^. Once mice began eating pellets, the turntable was gradually moved further away to encourage reaching for the pellet rather than licking. Once reaching began (within 2-5 days of initial exposure to the task), we started a training period of 10 consecutive days where mice performed 50 trials per day with a 30 s inter-trial interval. The position of the pellet for each mouse was adjusted by ∼0-2 mm within the first 1-2 days of training to match the preferred reaching position for each mouse. All mice undergoing this training paradigm achieved consistent expert performance during this training.

After 10-day training, mice began receiving the pellet delivered on an automated vertical post rather than the turntable (Janelia Experimental Technology) which improved visibility of the hand for capturing kinematics during neural recordings, as used previously^19^. Trials began every 30 s (except for inter-trial interval manipulations with 20 s and 40 s inter-trial intervals). Behavior was captured by three synchronized infrared cameras (PointGrey Flea3) at 500 Hz using BIAS (IO Rodeo) with event timing from WaveSurfer (Adam Taylor; Janelia Research Campus). Two cameras captured the front and side views and were calibrated with the Caltech Camera Calibration Toolbox. The third camera captured the pupil. Hand and digit kinematics (first and third digits) were tracked using the Animal Part Tracker (Kristin Branson; Janelia Research Campus). Post-processing was performed with custom MATLAB scripts to detect relevant time points: movement onset (first initiation of hand motion after the cue towards the pellet), maximum velocity (peak hand speed during the outward reach), hand open (peak of grip aperture), grab (onset of digit closure following the hand-open peak at the minimal hand-pellet distance), and supination (first sustained increase in hand rotation angle bringing the palm surface toward the mouth). Continuous kinematic features were also quantified: hand velocity, grip aperture (distance between first and third digits), and supination angle (hand rotation angle). Reaction time was measured as the time between trial cue and movement onset. Success was defined as successfully retrieving and consuming the food pellet. Action quality was defined as the mean Euclidean distance (mm) between each trial’s 3D hand position and the mean hand position of successful trials within that session, computed on 1 s of the hand trajectory (from −350 ms to +650 ms relative to movement onset). Lower values represent smaller trajectory deviations and correspond to higher action quality (i.e., reaches that are more similar to successful trials). For correlations with coupling and decoupling or firing rate changes, action quality was sign-inverted (multiplied by −1) so that positive correlations indicate better actions at higher coupling or firing rates.

### Multi-site neurophysiology

After 10-day training, we recorded neural activity as mice performed skilled reach-to-grasp actions. To do this, a second surgery was performed to place five craniotomies that allow acute insertions of recording electrodes and chronic implantation of a gold-plated ground pin (DigiKey). Kwik-cast (World Precision Instruments) was applied to temporarily cover the brain and skull after surgery and between recording days. The following day (or after 1-2 days of recovery), we performed simultaneous acute 4-shank Neuropixels 2.0^51^ recordings across five interconnected nodes of the motor network: frontal cortex (FC, coordinates: +1 to +2.8 mm AP, +0.6 to +1.0 mm ML), primary motor cortex (M1, coordinates: −0.1 to +0.7 mm AP, +0.4 to +2.2 mm ML), dorsolateral striatum (DLS, coordinates: −0.1 to +0.7 mm AP, +1.0 to +2.6 mm ML), motor thalamus (mTH; coordinates: −1.0 to −2.0 mm AP, +0.3 to +2.0 mm ML), and interposed nucleus of the deep cerebellar nuclei (DCN, coordinates: −5.75 to −6.5 mm AP, +1.1 to +2.6 mm ML). FC, M1, DLS, and mTH recordings were contralateral to the reaching hand, while DCN recording was ipsilateral to the reaching hand. The ground pin was placed contralateral to the reaching hand posterior to all craniotomies in that hemisphere (−4 mm AP, +2 mm ML). Four probes were used with M1/DLS recorded on the same probe. Shanks were coated in DiI stain (Thermo Fisher Scientific) to localize electrode track positions in post-hoc histology. Signals were acquired using SpikeGLX (Bill Karsh; Janelia Research Campus) at 30 kHz. Data simultaneously collected from multiple Neuropixels electrodes were aligned using sync channels in SpikeGLX. Spike sorting was performed using Kilosort 2.5^52^ with manual curation in Phy^53^ and automated quality control by Bombcell^54^. Units localized outside the anatomical coordinates listed above were excluded from analysis. Local field potentials (LFPs) were extracted using custom MATLAB scripts that low-pass filtered raw data at 300 Hz (fourth-order Butterworth) and downsampled to 1 kHz. Four channels (every 48th channel) were extracted from each shank of the 4-shank Neuropixels 2.0 probes (resulting in 16 total LFP channels per Neuropixels probe). Within each area, LFP channels were median-referenced. After recordings, mice were transcardially perfused with PBS followed by 4% paraformaldehyde. Brains were extracted, post-fixed overnight, and sectioned at ∼50μm using a vibratome. Sections were imaged to visualize DiI-labeled electrode tracks and confirm recording locations.

### Optogenetics

In *Slc32a1-ChR2(H134R*)*-EYFP* mice (*VGAT-ChR2; n =* 6 mice, 38 sessions), we placed two optical fibers targeting lobule simplex of the cerebellar cortex and M1. Fibers (200μm, NA 0.39, Thorlabs) were positioned on the skull surface at −6.3 mm AP +1.5 ML (lobule simplex) and +0.7 AP +2 ML (M1) and coupled to a 473 nm laser (Laserglow) via patch cords. Output power was measured at the fiber tip each session and adjusted to elicit reliable neuronal entrainment (range: 10-50 mW). During recordings, we simultaneously captured spikes in DCN and mTH to quantify phase-locking and cross-area spike-timing. Each session began with control (no-light) trials, followed by repeated cycles of 16 stimulation trials. On every stimulation trial, both sites received 4 s of 4 Hz sinusoidal light, ending 500 ms before the behavioral cue. The relative phase between cerebellar and cortical pulse trains varied across 16 offsets.

### Action modulation and timing

To quantify action-modulation in each neuron, we aligned spiking activity to movement onset on each trial (10 seconds before/after movement onset on each trial), binned activity into 2 ms bins (to match frame rate of cameras capturing behavior at 500 Hz), smoothed activity with a Gaussian kernel of 200 ms width, and then trial-averaged. Only successful trials were included in trial-averaging. We then z-scored each neuron’s trial-averaged movement-related activity using mean and standard deviation from a pre-movement baseline from −10 s to −2 s relative to movement onset. Units were classified as movement-modulated if the summed absolute value of the z-scored activity in a movement-aligned window (−500 to +1000 ms relative to movement onset) was more than twice the sum of a preceding pre-movement window (−2000 to −500 ms). Modulation sign was determined by whether the z-scored activity during the movement window showed a net increase (positively modulated) or decrease (negatively modulated) relative to pre-movement baseline. Modulation timing was defined as the first time bin within −500 to +100 ms relative to movement onset to cross a fixed z-threshold: +4 standard deviations for positively modulated units and −4 standard deviations for negatively modulated units.

### Action-informative and non-informative neuron classification

To identify neurons carrying movement information, we computed Shannon information using the MINT toolbox^55^ (direct method with naive bias correction) between each neuron’s firing rate (binned in 10 ms bins relative to movement onset and smoothed with a Gaussian kernel of 250 ms width) and kinematics (acceleration and grip aperture binned in 10 ms bins relative to movement onset, each feature computed separately). For each neuron, we generated a one-dimensional kinematic vector that concatenated binned kinematics from 0 to +500 ms from movement onset from all successful trials of a recording session (with magnitude discretized into five equipopulated levels). We then extracted the corresponding trial-concatenated binned and smoothed spiking activity at a range of time lags from kinematics (−500 to 0 ms between neural activity and kinematics in 10 ms steps), resulting in 50 total one-dimensional vectors of spiking activity (magnitude discretized into four equipopulated levels within each vector). Information was computed between each spiking and kinematic vector, providing an information profile over 50 neural-to-behavior time lags. To determine the significance of each time lag, we also computed a trial-shuffled distribution of information profiles (100 shuffles) by permuting trial order and recomputing information. We considered a time lag significant if it was part of ≥5 consecutive lags that exceeded the 95th percentile of this shuffled distribution. Neurons with ≥10 total significant time lags (out of 50) were designated action-informative, all others were non-informative. Neurons were considered action-informative if they contained significant information about either kinematic variable (acceleration or grip aperture). Information timing was defined as the earliest lag where a neuron first showed significant information about either kinematic variable.

### Principal components analysis (PCA)

To characterize population dynamics, we performed PCA on single-trial firing rates from neurons pooled across all recorded areas. For each neuron, single-trial firing rates were aligned to movement onset on each trial in 10 ms bins, then smoothed with a 250 ms Gaussian kernel. To quantify whether trajectories reflected kinematic variation (**Figure 1**), PCA was computed on the matrix of single-trial activity (time bins from concatenated trials as observations, neurons as features). PCA was performed in MATLAB using *pca* which mean-centers each neuron’s activity. On each trial, 5 s of spiking activity was included to compute PCA (2 s before movement onset to 3 s after movement onset). Trials were then grouped into quintiles based on grip aperture and acceleration profiles. For each trial, trajectory deviation was computed as the Euclidean distance between that trial’s neural trajectory (in the neural space defined by the top two PCs, from movement onset to 1 s after movement onset) and a template trajectory defined as the mean of the top quintile. We performed this analysis separately for three neuron populations: (1) action-informative modulated neurons, (2) non-informative modulated neurons, and (3) non-modulated neurons, to compare which population’s trajectories tracked kinematics. Sessions were included if each population contained at least 50 neurons. To quantify neural trajectory consistency (**Figures 3&4&6**), PCA was computed on a matrix of single-trial activity from action-informative neurons using a shorter period of spiking activity to specifically capture the consistency of neural trajectories during movement. On each trial 250 ms of spiking activity was included (0 to +250 ms relative to movement onset). Neural trajectory consistency was computed as the Pearson correlation between each trial’s trajectory (top two PCs) and the mean trajectory of successful trials. To assess whether principal component weights clustered by brain area, we examined the distribution of PCA weights for each area across the top two principal components.

### Local field potential (LFP) analysis

LFPs were extracted from Neuropixels recordings by low-pass filtering at 300 Hz (4th order Butterworth filter) and downsampling from 30 kHz to 1000 Hz. LFPs were median-referenced by subtracting the median signal across channels within each area. Spectral analysis was performed using the Chronux^56^ toolbox, with multitaper spectral estimation (time-bandwidth product = 3, 5 tapers). Power spectra were computed using MATLAB functions *mtspecgramc* and coherence using *cohgramc*, with a 1 s sliding window and 50 ms step size over a frequency range of 0-100 Hz. Trial-by-trial LFP power and coherence were computed between all pairs of recorded areas over a −10 to 0 s pre-movement window. Pre-movement changes in power and coherence were quantified as the difference between spectra averaged across late (−2 to 0 s) and early (−10 to −8 s) pre-movement periods. Statistical significance of spectral changes was assessed using Wilcoxon signed-rank tests with Bonferroni correction for multiple comparisons across frequencies. We focused on two frequencies for subsequent analysis: delta (δ, 4 Hz) and beta (β, 12 Hz). To assess single-unit phase locking, LFPs were bandpass filtered between either 2-8 Hz or 12-18 Hz using a zero-phase 2nd-order Butterworth filter, and instantaneous phase was extracted via the Hilbert transform. For each neuron, the phase of each spike occurring 2.5-5 s before movement onset was pooled across trials, and circular non-uniformity was tested using the Rayleigh test (CircStat toolbox^57^) to determine significant phase locking. To quantify the temporal relationship between pre-movement δ and β dynamics, we extracted δ and β time courses from trial-averaged coherence and power spectrograms. δ coherence was measured from the DCN-mTH interaction (the pair with the largest pre-movement δ coherence increase), and δ power was measured from DCN (the area with the largest pre-movement δ power increase). These δ traces were compared to β coherence or β power from all other area pairs or areas, restricting analyses to sessions in which δ increased and β decreased over the pre-movement interval. For each session, median coherence or power across channels were then mean-centered and their normalized cross-correlation was computed. The lag of the minimum cross-correlation value was taken as the timing of maximal anti-correlation. Session counts may vary across neuron-level and LFP analysis as not all sessions that yielded LFP also yielded isolated single units in every area.

### Cross-area coupling and decoupling

We defined coupling and decoupling based on the consistency of spike-timing relationships between cross-area neuron pairs. For each neuron on each trial, spiking activity was aligned to movement onset and binned at 10 ms intervals. The binned spike train was then smoothed using a Gaussian kernel with a 200 ms width. To remove trial-invariant components, the activity was median centered by subtracting the median across trials from each individual trial at each time point. For each pair of cross-area neurons recorded simultaneously, normalized cross-correlations were computed over a time window from −10 to +10 seconds relative to movement onset. A 500 ms sliding window with 250 ms steps was used to compute cross-correlations. For each time bin, spiking activity from the first neuron was paired with the activity of the second neuron delayed by an offset (−500 to +500 ms in 10 ms bins, resulting in 101 time lags compared). The first time bin included spiking activity of the first neuron from −9500 to −9000 ms relative to movement onset, while the last time bin included activity from +9000 to +9500 ms after movement onset, yielding a total of 75 time bins. Each cross-correlation value was computed by calculating the dot product of the activity of the two neurons (the sum of the element-wise product of their activity) and the norm of each neuron’s activity (sum of the squared values of their activity and taking the square root of the sum). The dot product and the norms were used to compute the normalized cross-correlation as cosine similarity by dividing the dot product by the product of the two norms. In cases where division by zero occurred (i.e., when either neuron had no activity), values were set to zero. This normalization was applied to minimize the impact of differences in firing rates between neuron pairs on our measure of their temporal relationship.

Pairwise directionality was assigned based on the lag of the peak in the trial-averaged cross-correlation of successful trials during the pre-movement period (−10 s to −0.5 s relative to movement onset), indicating which area’s spikes led or lagged the other. The peak lag was determined by first averaging the cross-correlation across pre-movement time bins, then identifying the lag with maximum magnitude. Neuron pairs were separated into two directional groups based on peak lag position: pairs with peak lags of −250 to −10 ms (first area leads) versus +10 to +250 ms (second area leads). For each neuron pair, we extracted the cross-correlation magnitude at this peak lag across all time bins, providing a dynamic readout of spike-timing consistency. Coupling and decoupling were defined by comparing cross-correlation magnitude between early (−10 to −6.5 s relative to movement onset) and late (−3.75 to −0.5 s) pre-movement periods. These time ranges account for the 500 ms sliding window used to compute each cross-correlation, as well as the peak lag offset (up to ±500 ms) between neuron pairs. Pairs showing an increase were classified as coupling, while pairs showing a decrease were classified as decoupling. For each recording session and directed area pair, we computed net coupling as the proportion of pairs that coupled minus the proportion that decoupled. Statistical significance was assessed using one-sample t-tests against zero with Bonferroni correction for multiple comparisons across all directed area pairs.

To relate coupling and decoupling dynamics to behavioral performance on individual trials, we computed trial-by-trial measures of coupling and decoupling. Neuron pairs were first classified as coupling or decoupling by comparing cross-correlation magnitude between early (−10 to −6.5 s relative to movement onset) and late (−3.75 to −0.5 s) pre-movement periods: pairs showing an increase were classified as coupling, while pairs showing a decrease were classified as decoupling. We then separately analyzed action-informative pairs that coupled and non-informative pairs that decoupled. For each trial, we computed the median cross-correlation magnitude across all pairs within each group. Trial-by-trial coupling magnitude was computed by z-scoring these values relative to the early pre-movement baseline (−10 to −6.5, mean and standard deviation computed across all trials and time points within this window), then averaging over a late pre-movement period (−6.25 to −0.5 s). The relationship between coupling/decoupling magnitude and action quality (see Reach-to-grasp task) and neural trajectory consistency (see Principal Components Analysis) was assessed using Spearman correlations across successful trials, with Bonferroni correction for multiple comparisons.

To relate coupling and decoupling dynamics to LFP signals, we computed trial-by-trial correlations between LFP coherence and coupling/decoupling magnitude. LFP coherence from area pairs with the largest pre-movement delta (δ, 4Hz) coherence increase (mTH-DCN) and beta (β, 12Hz) coherence decreases (M1-FC) were correlated to coupling/decoupling changes across all area pairs. For each trial, δ and β coherence were averaged across a pre-movement time window (−7 to −2 s relative to movement onset). Coupling and decoupling magnitude were computed as described above, averaged over the late pre-movement period (−6.25 to −0.5 s). The relationship between trial-by-trial LFP coherence and coupling/decoupling magnitude was assessed using Spearman correlations, computed separately for each directed area pair. Statistical significance was determined using Bonferroni correction for multiple comparisons (α = 0.05/20).

### Pre-movement firing rate analysis

To quantify pre-movement firing rate, spiking activity was aligned to movement onset and binned in 100 ms bins. For each session and area, pre-movement firing rate change was quantified as the proportion of neurons that increased or decreased firing rate from an early pre-movement period (−10 to −6.5 s relative to movement onset) to a late pre-movement period (−3.75 to −0.5 s). To relate the timing of pre-movement firing rate changes to changes in coupling and decoupling, firing rates were averaged across neurons within action-informative or non-informative populations on each trial. Correspondingly, action-informative coupling and non-informative decoupling were averaged across neuron pairs on each trial. Timing of change in either firing rate or coupling/decoupling was defined for each trial as the first pre-movement time bin in which the absolute magnitude exceeded double the median magnitude across the pre-movement period. Trial-level differences between firing rate onset and coupling/decoupling onset were then pooled across sessions and area pairs. To relate pre-movement firing rate to behavioral performance, single-trial firing rates were averaged within each neuron population (action-informative and non-informative) in each area, and normalized within session by subtracting the average firing rate from −10 to −6.5 s relative to movement onset. For each trial, the mean normalized firing rate from −3.75 to −0.5 s relative to movement onset was then correlated with action quality (see Reach-to-grasp task) using Spearman correlation, with Bonferroni correction across areas and metrics.

### Variable inter-trial interval manipulation

To test how disrupting coupling and decoupling impacted skilled action, we manipulated the inter-trial interval (ITI) in a subset of animals (*n* = 4) to initiate trials before typical coupling and decoupling dynamics emerged. After training at the standard 30 s ITI, we introduced short (20 s) and long (40 s) ITI trials on alternating recording days. On manipulation days, short or long ITI trials occurred once every six trials, alternating between short and long. On non-manipulation days, all trials used the standard 30 s ITI. Coupling and decoupling were analyzed separately for short, regular, and long ITI trials using the same methods described above. To ensure consistent pair classification across conditions, pairs were classified as coupling or decoupling based on their dynamics during regular ITI trials, then the same pairs were tracked across ITI conditions. Coupling magnitude was computed as the mean cross-correlation during the late pre-movement period (−3.75 to −0.5 s relative to movement onset). Statistical comparisons between ITI conditions were performed using two-sided Wilcoxon signed-rank tests.

### Pupillometry

Pupil diameter was extracted from high-speed videos using custom MATLAB scripts. Video frames were converted to grayscale and binarized using a manually adjusted threshold for each session. The resulting binary image was processed to isolate the pupil and pupil diameter was measured as the maximum pairwise distance between boundary points. Values were linearly interpolated across frames and resampled to match the kinematics sampling rate. Pupil diameter was z-scored within each session and aligned to movement onset. Pre-movement pupil change was computed as the difference between mean pupil diameter during a late (−2 s to −0.5 s) and early (−10 s to −8.5 s) pre-movement window. This pupil change was correlated with multiple neural features using Spearman correlations: mean delta (δ, 4 Hz) and beta (β, 12 Hz) LFP coherence across all recorded area pairs, and the pre-movement change in cross-correlation magnitude among action-informative neuron pairs that coupled and non-informative pairs that decoupled.

### Optogenetic manipulation analysis

To quantify neuronal entrainment to laser stimulation, we computed the phase of each spike relative to the 4 Hz laser waveform during the 4 s stimulation period. Instantaneous phase of the laser stimulation was extracted using the Hilbert transform. For each neuron, spikes occurring during the stimulation period were pooled across all laser trials relative to the laser at the corresponding stimulation site (motor cortex laser for mTH neurons, cerebellar laser for DCN neurons). Significant phase locking was assessed using the Rayleigh test for circular non-uniformity (P<0.05, Bonferroni-corrected). The preferred phase was defined as the center of the histogram bin (π/16 width) containing the most spikes. To assess how phase-offset stimulation affected cross-area spike timing, we computed cross-correlations between DCN and mTH neuron pairs using the same methods described above. Cross-correlations were computed separately for each of the 16 laser trial types and for non-laser control trials. The similarity between laser-induced and control cross-correlations was quantified as the Pearson correlation between cross-correlation profiles for each laser trial type group and the profile from successful non-laser control trials for cross-area lags of ±100 ms. The effect of phase-offset stimulation on behavior and neural activity was assessed by comparing action quality and LFP power across laser trial types grouped using a sliding window of 3 consecutive types. Action quality was quantified as trajectory deviation (see Reach-to-grasp task). Beta (β, 12 Hz) LFP power in mTH was computed during the pre-movement period. For each measure, values were expressed relative to the mean across all 16 laser trial types within each session. Statistical significance was assessed using Wilcoxon signed-rank tests against zero with Bonferroni correction for multiple comparisons across trial type groups.

